# Prey size spectra and predator to prey size ratios of southern ocean salps

**DOI:** 10.1101/2022.02.16.480784

**Authors:** Christian K. Fender, Moira Décima, Andres Gutiérrez-Rodríguez, Karen E. Selph, Natalia Yingling, Michael R. Stukel

## Abstract

Salp grazing is important in shaping planktonic food-web structure. However, little is known about the size ranges of their prey in the field or how grazing impacts size structure. This study investigated the feeding habits of 7 different species of salps, representing a variety of sizes and life stages across subtropical and subantarctic waters east of New Zealand. Scanning electron microscopy was used to examine the gut contents of 58 salps, which were then compared to water column plankton communities characterized via epifluorescence microscopy, FlowCam, and flow cytometry. While most of the gut contents resembled ambient waters, substantial differences were found amongst some co-occurring species, such as increased retention of submicron bacteria amongst smaller salps like *Thalia democratica*. We found that even for those salps capable of feeding on bacteria efficiently, nanoplankton and small microplankton still made up the majority of gut biomass. Larger microplankton were rarer in the guts than in the water column, potentially suggesting an upper size-threshold in addition to the lower size-threshold that has been the focus of most previous work. Salp carbon-weighted predator to prey size ratios were variable, with the majority falling between 1,000:1 and 10,000:1 depending largely on the size of the salp. Taken together our results indicate that despite being able to feed on submicron particles, picoplankton make up at most 26.4% (mean = 6.4%) of salp gut carbon and are relatively unimportant to the energetics of most salps in this region compared to nanoplankton such as small dinoflagellates and diatoms.

## 1. Introduction

Salps are a group of large (∼1 – 30 cm as adults), suspension-feeding gelatinous zooplankton that filter water through a fine mucous mesh at hourly rates of more than 1000 times the organism’s biovolume (Madin et al. 2006). With population doubling times as fast as 8 hours due to their dual staged life history composed of both solitary and aggregate forms (Alldredge and Madin 1982; Deibel and Lowen 2012), salps can form swarms of up to 1000 individuals m^-3^ covering up to 9065 km^2^ in response to favorable conditions (Anderson 1998; Berner 1967). Salps can have clearance rates >100,000 mL ind^-1^ day^-1^ (Madin and Kremer 1995; Perissinotto and Pakhamov 1998; Sutherland et al. 2010), equivalent to the clearance rate of 450 copepods (Harbison and Gilmer 1976), allowing these salp blooms to clear water columns of prey so quickly that they can prematurely end spring diatom blooms before surface nutrients are depleted (Bathmann 1988). Despite a growing appreciation for the impact salps may have on ecosystem structure and biogeochemical cycles, there are still gaps in our understanding of their feeding ecology such as the distribution of sizes over which they feed. The ratio of a predator’s size to that of its prey has long been linked to the total length of an ecosystem’s food web, where large predator:prey size ratios (hereafter PPSR) result in fewer trophic levels and more efficient transport of primary production to large taxa rather than remineralization (Lindeman 1942; Sheldon et al. 1977; Sherr and Sherr 1988). While it is less frequently recognized, the range of prey sizes a predator can feed upon has similar impacts on ecosystem structure. More generalist predators display higher standard deviations in prey size (hereafter SD_PPSR_) resulting in ecosystems with lower connectance, smaller phytoplankton, and less diversity in phytoplankton size classes (Fuchs and Franks 2010). In short, the PPSR and SD_PPSR_ of the dominant predators in an ecosystem play a strong role in determining its trophic structure, and only a handful of studies have quantitatively investigated the size of salps’ prey in the field (Dadon-Pilosof et al. 2019; Madin 1974; Madin and Purcell 1992; Vargas and Madin 2004).

It has long been accepted that the relationship between predator and prey size has a strong impact on the ecology of an ecosystem (Sheldon et al. 1977; Jennings et al. 2002) and that PPSR broadly varies between zooplankton of different feeding modes (Hansen et al. 1994). For instance, while most zooplankton tend to feed with PPSRs between 10:1 and 100:1 (Hansen et al. 1994; Fuchs & Franks 2010), raptorial protists often feed with PPSRs of closer to 3:1 and both pallium-feeding dinoflagellates and some siphonophores can feed at a ∼1:1 PPSR (Purcell et al. 1981; Fenchel 1987; Sherr et al. 1991; Naustvoll 2000; Sherr & Sherr 2002). At the other end of the spectrum, for filter-feeding plankton it is common to observe anywhere from a 5:1 to 100:1 ratio between an organism’s size and that of its prey (Hansen et al. 1994; Conley et al. 2018). These variations are the product of physiological differences between raptorial and filter feeders that lead to significant differences in the portion of available prey biomass that is utilized (Figure 1). For example, raptorial crustaceans can be expected to exhibit a preferred or optimal prey size whereas the size distribution of prey for filter feeders is more broadly determined by what is available in the water column. Filter feeders also tend to have higher feeding efficiencies for small particles compared to raptorial feeders of similar size, differentiating the niche space such that filter feeders are often in competition with organisms far smaller than themselves (Stukel et al. 2021). Depending on the size of the filter feeder, this is manifested in substantially higher PPSRs. High feeding efficiencies for a broader range of prey sizes likewise result in a higher SD_PPSR_.

**Figure 1.**
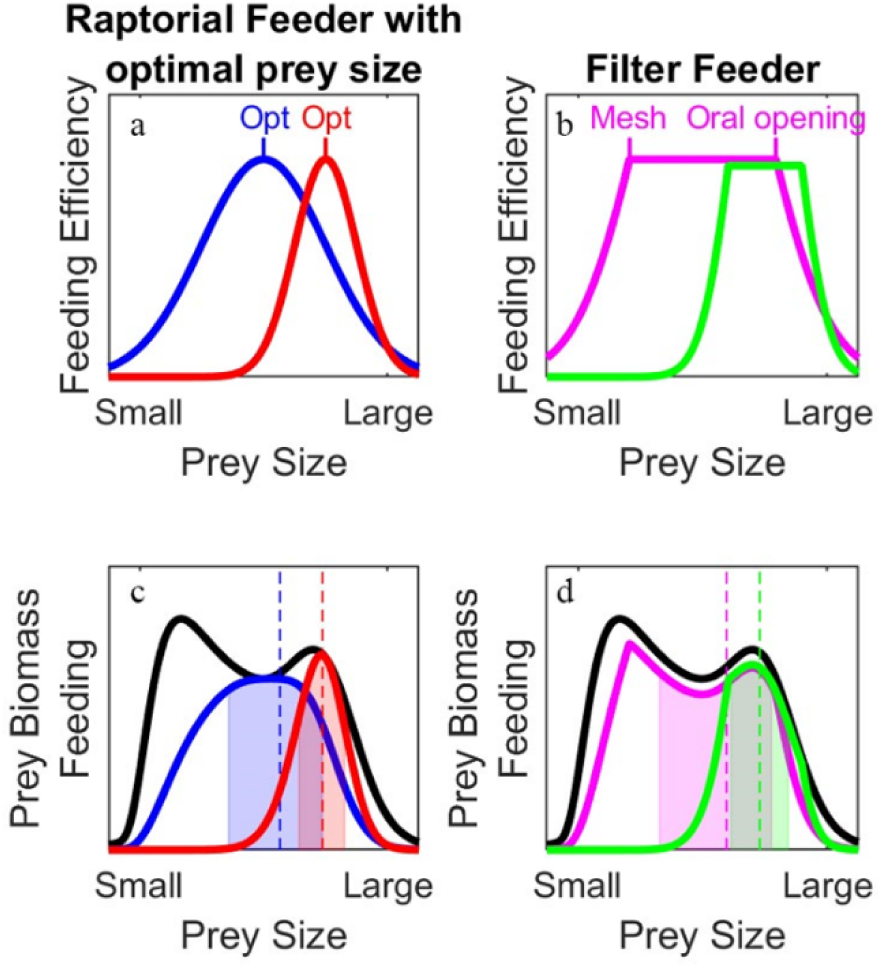
Feeding efficiencies for two theoretical raptorial feeders with an optimal prey size (a) and for two theoretical filter feeders whose feeding efficiency as a function of prey size is determined by the mesh size of their filters and the size of their oral opening. (b) Blue and purple lines represent organisms with higher SD_PPSR_ than the red and green organisms. Hypothetical prey biomass as a function of size is shown in black in (c) and (d) along with feeding rates as a function of size, which is equal to prey biomass times feeding efficiency for hypothetical raptorial (c) and filter (d) feeders. Vertical dashed lines show the carbon-weighted mean prey size (i.e., the prey size for which half of prey biomass consumed was greater than that size and half was less than that size), while prey sizes within one standard deviation of the mean prey size are shown in the shaded colors. PPSR for a given salp was computed as the salp’s length divided by its carbon-weighted mean prey size.

As an extreme example of the efficiency of filter feeders, pelagic tunicates (salps, doliolids, pyrosomes, and appendicularians) often feed with PPSRs >1,000:1 (Madin 1974; Crocker et al. 1991; Madin and Purcell 1992; Vargas and Madin 2004; Katechakis et al. 2004). This is accomplished through the production of a fine mucous mesh secreted by the endostyle and passed down to the esophagus through the action of cilia lining the gill bar which passes through the middle of the organism (Madin et al. 1974; Sutherland et al. 2010). As the salp compresses the muscle bands lining its thick outer test, filtered water is forced out of the posterior aperture with fresh water replacing it through the anterior aperture as the muscles relax. This creates a form of jet propulsion the organism utilizes for locomotion, and as the water is forced through the interior cavity it is passed through the feeding filter such that food particles are entrained. This allows salps to feed directly on micron-sized picoplankton, but also consume ∼1-mm organisms including copepod nauplii, radiolarians, foraminifera (Madin 1974), and ostracods (Décima et al. 2019).

Consequently, salps have the potential to simultaneously act as competitors to and grazers of a wide variety of phagotrophic protists (Stukel et al. 2021), the group responsible for the majority of phytoplankton grazing (Calbet and Landry 2004). Furthermore, as salps themselves are part of the diet of at least 202 marine species, salps may not act as a net carbon sink as classically believed but instead increase trophic transfer and vertical export efficiencies (Michaels and Silver 1988; Henschke et al. 2016). Their presence could thus support economically important species like mackerel (Nishimura 1958), bluefin tuna (Cardona et al. 2012), anchovies (Mianzan et al. 2001), and other demersal fishes (Horn et al. 2011; Forman et al. 2016). However, the physiological ability to feed at PPSRs up to 10,000:1 does not necessarily mean that this is the range over which most salps feed. As non-selective filter feeders, it is likely that this value is largely dependent on the phytoplankton community present. Differing retention efficiencies for small particles have also been reported for a variety of salp species based on removal studies in deck-board incubators (Kremer and Madin 1992; Vargas and Madin 2004; Stukel et al. 2021), though these techniques often struggle to accurately resolve very small or very large prey. The size of the salp itself likely plays some role as well, as there is evidence that small salps may retain smaller particles more efficiently than larger individuals even of the same species (Harbison and McAlister 1979; Kremer and Madin 1992; Stukel et al. 2021), and that the size of the oral opening (i.e. the esophageal opening in salps; Madin 1974) will limit consumption of larger-sized phytoplankton cells and/or diatom chains (Figure 1b). Whether feeding differs intrinsically between species or life stages or if it is purely a function of size is also unknown.

Although traditional microscopic analyses of salp stomach contents have long supported their non-selective nature (Silver 1975; Vargas and Madin 2004; Tanimura et al. 2008), these methods are limited in terms of image resolution such that only large, hard-bodied plankton can be easily identified. Modern tracer and genetic analyses have also been applied to gut contents to determine relative proportions of different types of prey (von Harbou et al. 2011; Metfies et al. 2014; Pauli et al. 2021; Thompson et al. 2023) and are beginning to challenge the existing paradigm of non-selectivity, as these studies more often find differences between prey types available and those found in the guts. However, these methods do not allow for determining prey size. A handful of studies have also attempted to directly assess salp gut contents via scanning electron microscopy (SEM), which can resolve particles from the submicron to millimeter size range. Unfortunately, preparatory methods vary, and the focus has mostly been on larger, more easily identifiable prey items like diatoms and thecate dinoflagellates (Madin and Purcell 1992; von Harbou et al. 2011; Ahmad-Ishak 2017). This, along with observations of the mesh spacing in their mucous filters, have led to the commonly accepted view that most salps predominantly feed on particles >1-2 µm in diameter due to the inefficiency with which smaller particles are retained (Harbison and McAlister 1979; Kremer and Madin 1992; Madin and Purcell 1992). Some studies leveraging numerical simulations as well as removal experiments using beads of known size have recently challenged this, suggesting that direct interception of submicron particles by the mucous fibers may be a more important source of nutrition than originally believed (Sutherland et al. 2010). Even if retained less efficiently than larger particles, submicron phytoplankton like cyanobacteria are present at far higher concentrations than larger taxa such that feeding on them may still satisfy a large portion of a salp’s carbon quota (Sutherland et al. 2010; Dadon-Pilosof et al. 2019).

In this study, we sought to compare the size composition of prey items in salps’ stomachs to that of the water column, determine if the resulting size spectra are a function of salp species, size class, or life stage, and then quantify how these demographics impact the ecosystem as a whole through the PPSR, SD_PPSR_, and retention efficiencies of salps. We utilized gut content SEM analysis of 58 individuals comprised of both life stages of 7 different salp species ranging from 8-163 mm to quantify their diets and compare these diets to water column plankton populations characterized by flow cytometry, epifluorescence microscopy, and FlowCam imaging. To our knowledge, this is the most diverse collection of salp diets sampled to date in a single study.

## 2. Materials and Methods

### 2.1 Field collection

Samples were collected over Aotearoa New Zealand’s Chatham Rise during October 21 - November 21 2018 as part of the Salp Particle Export and Ocean Production (SalpPOOP) study, which was designed to investigate how salps affect the ecology and biogeochemistry of the region (Décima et al. 2023). The Chatham Rise is notable because it sits within the Subtropical Front of the southwest Pacific, which defines the boundary between warm, salty and low-nitrate subtropical waters and cold, fresh, nitrate-rich, and iron-poor subantarctic waters (Zentara and Kamykowski 1981; Heath 1985; Sokolov & Rintoul 2009;). This creates a dynamic frontal zone with high mixing that supports high phytoplankton biomass and productivity (Currie and Hunter 1998) as well as, anecdotally, periodic summertime salp blooms. The cruise consisted of 5 quasi-Lagrangian experiments (hereafter “cycles”) lasting 4-8 days each, with only 4 cycles (Cycle 1-4) showing significant salp presence (Figure 2). Cycle locations were chosen based on net tows through regions with high potential for salp presence according to previous observations on habitat distributions of salp-eating demersal fish (Forman et al. 2016; Horn et al. 2011).

**Figure 2.**
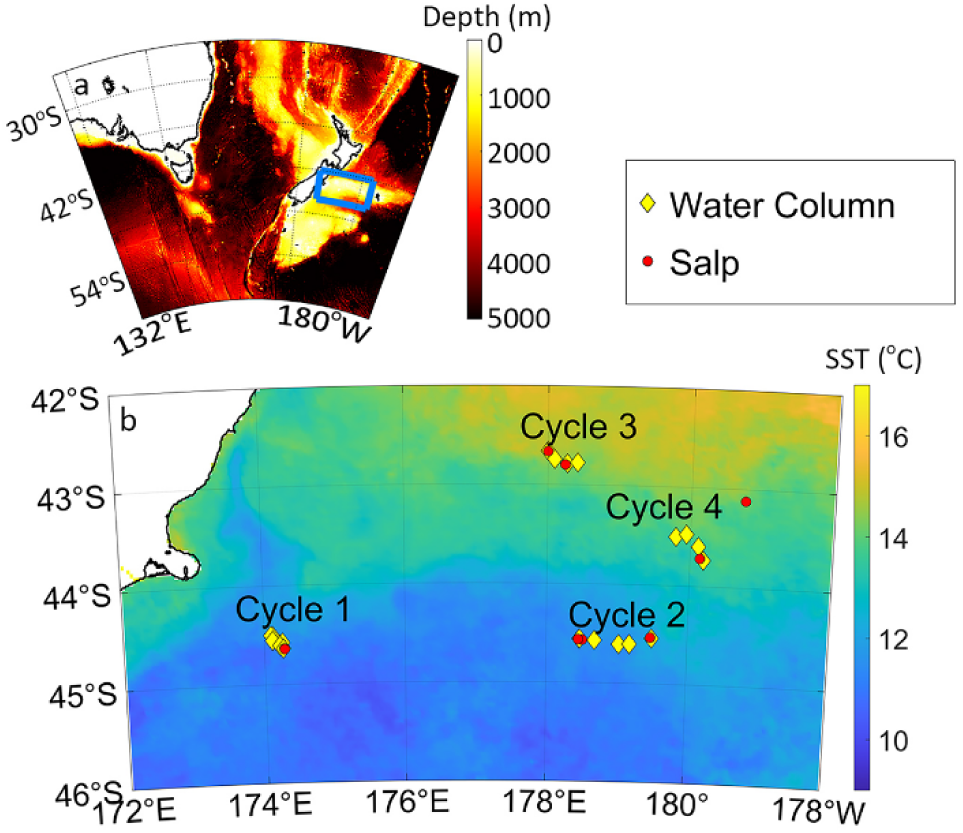
a) TAN1810 cruise study area (blue square) located east of Aotearoa New Zealand with color denoting bottom depth. b) Magnification of study region with color denoting sea surface temperature. Yellow diamonds represent CTD deployments to determine water column properties and red circles represent the locations of salp collections for SEM via Bongo or ring net tow.

At the beginning of each cycle, we deployed a surface-tethered drifting array to track the chosen water parcel (Landry et al. 2009). Daily dawn and noon deployments of a 24-bottle CTD-Niskin rosette were made to collect water used in bottle-incubation experiments as well as water column profiling. To characterize the prey community, we analyzed 4 types of samples collected from Niskin bottles: (1) 250 mL subsamples from the base of the mixed layer and the deep chlorophyll max (DCM) were concentrated by gravity filtration to 10 mL over a 2 µm 47 mm filter, and 2 mL of this concentrate was imaged using a FlowCam model VS-IV’s 10X objective lens to quantify the larger (>4 µm) phytoplankton (Sieracki et al. 1998); (2) an Accuri C6 Plus flow cytometer was used at sea to determine the abundance of *Synechococcus* and phototrophic eukaryotes collected at 6 depths spanning the euphotic zone with cell size estimated using the forward light scatter of polystyrene beads of known size (Stukel et al. 2021); (3) additional preserved samples were collected from the same casts (same 6 depths) and analyzed on a Beckman Coulter CytoFLEX S flow cytometer to enumerate heterotrophic bacteria and *Prochlorococcus* (Selph 2021); (4) preserved and stained microscopy samples were collected from 6 depths per cast and later analyzed on an epifluorescence microscope where cell biomass and abundance were calculated using ImageJ (Taylor et al. 2015). Each of these four methods allowed for the enumeration of overlapping parts of the phytoplankton size spectra, namely 0.4-50 µm for flow cytometry, 2-200 µm for epifluorescence microscopy, and 4-300 µm for FlowCam. In total, 21 CTD casts were sampled over the course of the 4 cycles from which salps were collected.

To determine salp biomass and demographic structure, we conducted twice daily oblique Bongo tows down to 200 m as well as MOCNESS tows to deeper depths of up to 2600 m twice per cycle. Ring net surface tows with a 30 L non-filtering cod-end were also conducted daily to collect additional salps. Once onboard, salps from ring net and Bongo tows were identified to the species level, sorted by life stage (i.e., solitary or aggregate), measured, and sexed (Foxton et al. 1966; Lüskow et al. 2020). Triplicate representative samples for each salp species from a total of 10 of these casts were preserved in 5% formalin < 30 minutes after collection the first time each species was encountered. While this means there was potential for prey digestion between the time the salp was caught and when preservation occurred, processing time was always substantially less than the estimated gut pigment turnover time of 2.8 hours for even the smallest salp caught (von Harbou et al. 2011). Whenever possible these preserved samples included a distribution of size classes and both solitary and aggregate life stages.

### 2.2 Scanning electron microscopy

In total, 58 salps representing the species *Salpa thompsoni*, *Pegea confoederata*, *Thalia democratica*, *Soestia zonaria, Thetys vagina*, *Salpa fusiformis*, and *Ihlea magalhanica,* including both solitary and aggregate stages of the first four and a distribution of sizes for the first two, were collected. Once ashore, SEM samples were prepared from each preserved organism by excising guts under a HEPA-filter equipped laminar flow exhaust hood using acid-cleaned plastic dissection equipment to minimize contamination. Guts were then placed in either 15 or 50 mL plastic Falcon tubes with a small volume of brine, lacerated, and then vortex mixed for two minutes to release gut contents into solution while minimizing damage to the more fragile phytoplankton (Jung et al. 2010; von Harbou et al. 2011; Ahmad-Ishak 2017). An aliquot of this solution was then filtered onto a 0.2 μm Nuclepore filter, followed by six rinses of decreasing salinity in 5 ppt increments for a minimum of 5 minutes each with the final MilliQ water rinse performed twice. This was immediately followed by a dehydration series of increasing ratios of Ethanol:MilliQ to purge water from the sample, with the final 100% anhydrous ethanol step again performed twice. Finally, a substitution series of increasing ratios of the chemical drying agent hexamethyldisilazane (HMDS):anhydrous ethanol was conducted with each step lasting a minimum of 10 minutes, with the final HMDS step being allowed to air dry. We chose this chemical drying agent over more traditional critical point drying both to minimize changes in cell size as well as maintain material on the filter (Jung et al. 2010). Each step was conducted under either a light vacuum or gravity filtration depending on material concentration to minimize loss between treatments. Similar procedures have proven successful in the preparation of both delicate dinoflagellates (Botes et al. 2002; Jung et al. 2010) and bacteria (Koon et al. 2019).

The dried filter was then affixed to an aluminum SEM stub using carbon conductive adhesive tabs and further grounded with a thin piece of carbon conductive tape touching the edge of the filter and the bottom of the stub. Samples were then sputter coated with 10 nm iridium and visualized using an FEI Nova 400 NanoSEM set to an accelerating voltage of 10 kV. Twenty random regions of each filter were imaged at 3 different magnifications: ∼500x (corresponding to a 500 µm by 500 µm imaging area), ∼2,500x, and ∼12,000x to target microplankton (20-200 µm), nanoplankton (2-20 µm), and picoplankton (<2 µm), respectively. Since sufficient structural detail could not be observed to definitively identify very small particles, spherical particles within the size range of ∼0.4-1.5 µm and resembling control images from lab cultures of *Prochlorococcus sp.* and *Synechococcus sp.* are instead referred to as bacteria-like particles. We also note that while formalin preservation can lead to cell shrinkage (Choi and Stoecker 1989; Zinabu and Bott 2000), we assume that the shrinkage would be roughly proportional amongst all cells and therefore does not impact our relative contributions of different groups to the nutrition of the salps. Furthermore, we assume that this shrinkage will have an effect on the dimensions of most phytoplankton taxa that is within our margin of error with the exception of ciliates, for which we apply a carbon conversion specifically for formalin fixed cells, and nanoflagellates. Studies have shown ∼40% decreases in cell volume for cultured flagellates due to formaldehyde fixation (Choi and Stoecker 1989; Zinabu and Bott 2000), which translates to a 10-20% decrease in ESD. This roughly corresponds with the 20% decrease in total length of salps preserved in 5% formalin noted by Madin et al. (2006) which will have a small impact on the absolute lengths of both predator and prey reported here (although this small difference is within our margin of error) but is unlikely to affect the PPSR.

### 2.3 Image processing

Particles in SEM and epifluorescence microscopy images were manually outlined using ImageJ (v. 1.52a or 1.53c) to extract the maximum feret length (L), minimum feret length, and area (A). These measurements were used to estimate equivalent spherical diameter (ESD), biovolume (BV), and carbon biomass assuming a prolate spheroid. To avoid overestimating the size of irregularly shaped particles, we calculated width (W) for a prolate spheroid of measured A and L such that:

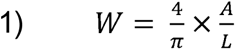

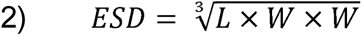

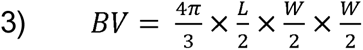

Note that because we estimated the three-dimensional size of particles using a two-dimensional image, we assume the height of each particle to be equal to its width. To account for the ∼50% flattening of height caused by filtration (Taylor et al. 2011), for all compressible particle types we instead assume height to be equivalent to half of the width. The biomass of formalin-preserved ciliates was estimated as 0.14 pg C µm^-3^ (Putt and Stoecker 1989) while rhizarians were 0.001 pg C μm^-3^ (Stukel et al. 2018). The biomass of diatoms was estimated allometrically as 0.288*BV^0.811^ while other protists and unidentified particles were estimated using 0.216*BV^0.939^ (Menden-Deuer and Lessard 2000). Because we could not differentiate between types of prokaryotes in the SEM images, we calculated a single average biomass conversion for all bacteria-like particles in the salp guts using published allometric relationships weighted by the ratio of each of the key bacterial groups to each other in the water column from flow cytometry data for a given cycle (see Supplementary Methods).

FlowCam image analyses were conducted using FlowCam’s dedicated classification software VisualSpreadsheet (v. 4.18.5). First, duplicate images resulting from parabolic flow within the flow cell were manually removed. The particles within the remaining images were then classified based on the quality with which VisualSpreadsheet detected their outlines. For particles where the outline appeared to provide good estimates of L and W, size was calculated for a prolate spheroid as above using Equations 1-3 with the exception that W was left as the directly measured minimum feret length and no correction for flattening was applied. While this is likely less accurate than utilizing particle area as was done for the SEM images, we frequently noted that particle outlines possessed large holes not representative of overall shape such that area was severely underestimated. For particles where the quality of automatic outline detection was poor, alternative methods used to determine size can be found in the Supplementary Materials. Biomass was calculated as described above for the salp gut contents with the exception of a non-formalin preserved conversion for ciliates of 0.19 pg C µm^-3^ (Putt and Stoecker 1989). For epifluorescence microscopy, biomass was similarly calculated using either the allometric equations from Menden-Deuer and Lessard (2000) for diatoms or for other protists. Carbon content for cells detected via flow cytometry were also calculated based on biovolume using the relationship for protists from Menden-Deuer and Lessard (2000), with the exception of *Prochlorococcus* (36 fg cell^-1^; Buitenhuis et al. 2012) and heterotrophic bacteria (11 fg cell^-1^; Garrison et al. 2000) for which we used fixed carbon conversions.

### 2.4 Size spectra and feeding distribution parameter calculations

For each cycle, the prey items in the salp guts were separated into size bins and their abundance was averaged for each salp species. This abundance was then divided by the size bin width to calculate normalized abundance size spectra (NASS). NASS were similarly produced for each of the methods used to assay the prey field (FlowCam, flow cytometry, and epifluorescence microscopy samples) at each depth and vertically integrated over the euphotic zone. We also present a single composite water column spectrum representing the geometric mean of each method over the size ranges where it was relevant. Carbon biomass estimates for both salp and water column measurements were then used in the same way to compute normalized biomass size spectra (NBSS). 95% confidence intervals were calculated using Monte Carlo random resampling (see Supplementary Methods for details).

During SEM analyses, we noted a surprising lack of diatom chains compared to the water column. We also noted 8% of all particles identified appeared to be broken (less than ¾ of their assumed true length) of which half were also chain-forming diatoms. To avoid biasing our salp gut size spectra analysis towards smaller particles when comparing against that of the water column where intact diatom chains were still present, we corrected our dataset for chain separation and cell breakage. In short, we utilized a similar Monte Carlo random resampling scheme to reassign abundances and biovolumes for all broken particles as well as chain-forming taxa in accordance with their size distributions in the water column and estimate any additional introduced uncertainty (See Supplementary Methods 2.4.2). While this solution does correct for both particle breakage and chain disruption that may have occurred between the time the particle was entrained and when it was imaged, we do note it may also obscure real differences between the salp gut plankton population sizes and those of the water column. For this reason, while all figures in the main text include broken particles, we also frequently provide alternative versions excluding them (but still correcting for chain disruption in diatoms) in the Supplement section. It is worth noting that inclusion of these particles yields little to no impact on any of our conclusions.

The carbon-weighted mean prey size of each salp was calculated by sorting all particles in a gut from smallest to largest and finding the prey size at which 50% of the total biomass in a salp gut was achieved (Supplementary Figure 2). The total length of a given salp was then divided by this value to estimate the carbon-weighted predator:prey size ratio (PPSR). We also calculated the SD_PPSR_ by finding the particle size at which ±1 standard deviation, or 15.9% and 84.1%, of a given salps total gut carbon content was met and then dividing the difference of the log transformed values by 2. The results are thus feeding distribution values that are more representative of the energetic value of the salps’ diets.

The degree to which each salp can efficiently retain picoplanktonic prey is determined by the mesh spacing of their feeding filter, which varies between species (Bone et al 2003). We estimated retention efficiency (hereafter RE) by dividing the prey NASS consumed by each salp species by that occurring in the water column for each size class and dividing the result by the average clearance rate for the 8-32 µm particles as this is the size range expected to be retained with 100% efficiency. We chose the 8-32 µm size range because particles with ESDs in the 4 – 8 µm size range will sometimes possess aspect ratios allowing them to pass through the feeding mesh and our results show that particles larger than this were frequently absent from salp guts.

## 3. Results

### 3.1 Salp vs. ambient phytoplankton community comparisons

Cycle 1, which occurred closest to the coast of the 4 cycles and was most strongly influenced by the Southland Current, a coastal expression of the Subtropical Front (Sutton 2003), was primarily dominated by *Salpa thompsoni* with smaller numbers of *Soestia zonaria* and *Thetys vagina* (Supplementary Table 1). Its phytoplankton community had the largest contribution of microplankton (20-200 µm) to the water column (66.8% of total vertically integrated plankton carbon, Figure 3) and it was also the only cycle where phytoplankton larger than 87 µm were detected. Nanoplankton (2-20 µm) and picoplankton (<2 µm) contributed 20.9% and 12.3% of the carbon, respectively. Qualitative analysis of FlowCam images from Cycle 1 suggests the microplankton were mostly centric and pennate diatoms. Flow cytometry data showed that *Synechococcus*, picoeukaryotes, and heterotrophic bacteria were present in all 4 cycles, while *Prochlorococcus* sp. was only present during Cycle 2.

**Figure 3.**
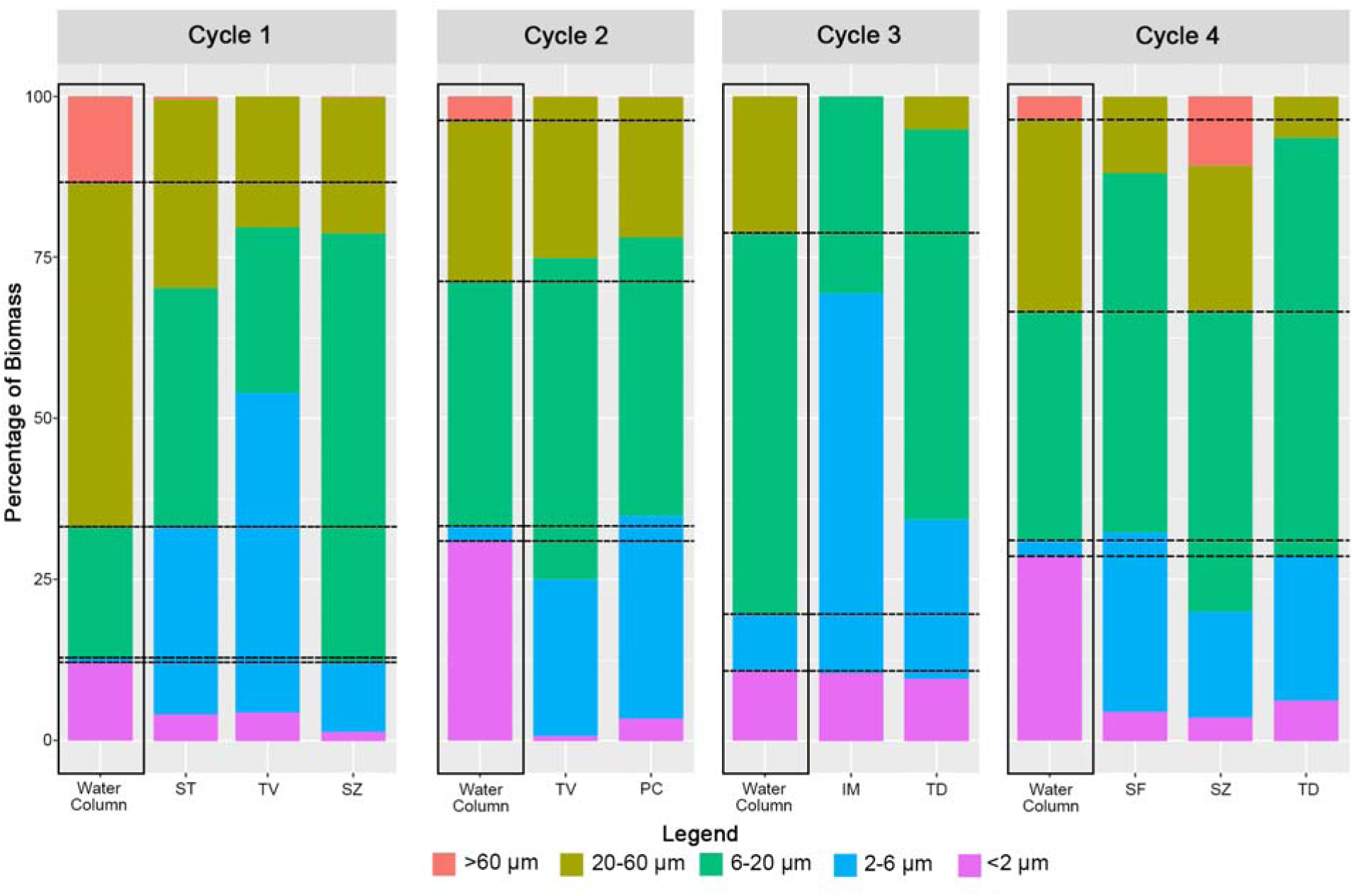
Contribution of particles of various size classes to the total biomass present in the water column (outlined by black boxes) compared to that of the guts of each of the 58 salps sampled averaged by species (as indicated on the x-axis) per cycle. WC = water column, SF = *Salpa fusiformis*, ST = *Salpa thompsoni*, TV = *Thetys vagina*, SZ = *Soestia zonaria*, PC = *Pegea confoederata*, IM = *Ihlea magalhanica*, TD = *Thalia democratica*. Horizontal dashed lines extend the WC demarcations for easier comparison to salp species.

Cycle 2 was farther from the coast and, similarly to Cycle 1, supported a salp community of primarily *S. thompsoni* along with subpopulations of *T. vagina*, *S. zonaria*, *Pegea confoederata*, and *Ihlea maghalanica* with a phytoplankton community characterized by a smaller proportion of microplankton (28.8% of total carbon) dominated by diatom assemblages that may have been advected off the coast. The contribution of nano-and picoplankton, however, was approximately twice that of Cycle 1 at 40.2% and 30.9%, respectively.

Cycle 4, which was conducted approximately 1° north of Cycle 2 in waters 3.1°cC warmer (Figure 2), had very similar size contributions as Cycle 2 albeit with nearly all of the microplankton portion made up by large dinoflagellates rather than diatoms. This supported a salp community that still had a high abundance of *S. thompsoni* but was also codominated by *Thalia democratica* along with background levels of a variety of other species.

Cycle 3 exhibited no clear salp blooms with only low to moderate abundances of *S. zonaria*, *I. maghalanica*, *S. fusiformis*, and *T. democratica* (although *T. democratica* was absent from MOCNESS tows and only found in ring net/bongo net catch). Only 21.2% of its total carbon was in the microplankton size range, with no >60 µm cells observed. Instead, 67.9% was comprised of nanoplankton, twice that of Cycles 2 and 4 and four times that of Cycle 1. The contribution of picoplankton was comparable to Cycle 1 at 11.0%.

Microplankton within salp guts were broadly representative of the taxa present in the water column with some notable exceptions elaborated on in the Discussion. When considering the contribution of various size classes to the gut content of all 58 salps, picoplankton made up between 0.5% and 23.2% of total biomass (Figure 3) with a mean and standard deviation of 4.8±4.5%. This is substantially lower than the cycle averaged mean contribution in the water column of 20.7±10.5%. The most abundant objects observed in the guts of all salps regardless of size or species were unidentifiable, smooth, spherical particles (hereafter referred to as “white spheres”) lacking any other notable morphological features and generally ∼2-7 µm in diameter but ranging from 1.5-20.5 µm (Supplementary Figure 3). Given the broad range in size and numerical abundance (they made up 78.3% of all identified nano-sized particles), it is likely that this category comprises several unrelated taxonomic groups that had been partially digested, potentially including prymnesiophytes, pelagophytes, prasinophytes, cysts (diatoms or dinoflagellates), and/or other nanoflagellate taxa. These, along with large numbers of similarly sized dinoflagellates, contributed to high concentrations of nanoplankton within the gut contents (range = 30.6% - 99.2% of total gut biomass, with a mean of 75.6±16.5%). This is overall substantially higher than what was observed in the water column as nanoplankton made up only 41.7±19.4% of the biomass on average. Microplankton were generally much rarer in the salp guts (range = 0% - 67.7% of biomass, mean = 19.6±17.8%) in contrast to the available prey in the water column (water column microplankton = 37.6±20.1% of biomass).

We also compared the average size spectra of prey for each salp species to that of the water columns of each of the four cycles from which samples were obtained (Figure 4). Since the volume filtered could not be determined for the salps caught, the units for the normalized abundance size spectra (NASS) of the prey and normalized biomass size spectra (NBSS) differ between salp and ambient measurements such that only the shape of the spectra can be compared. This shape compares the relative importance of small to large particles with steeper (i.e., more negative) spectra indicating more small particles relative to large ones and vice versa. In general, most of the salp ingested-prey spectra match up well to both the water column NASS and NBSS as would be expected for non-selective filter feeders. However, every salp species shows the same sharp decline in the abundance of consumed submicron particles, which supports the assumption of inefficient capture of particles <1 µm. There are several other notable departures from the ambient spectra as well. For example, in *Thalia democratica* of Cycle 3 the NBSS spectra is much flatter than that of the flow cytometry. While this trend is less apparent for *T. democratica* observed in Cycle 4, it could suggest an increased importance of small particles for this species relative to the others. In contrast, the NASS and NBSS spectra for *Ihlea magalhanica* is far more peaked at 2-4 µm compared to ambient; rather than suggesting difficulty in retaining small particles, this shape is likely due to the far higher number of white spheres relative to any smaller or larger particles observed in their guts. *Salpa thompsoni* and *Soestia zonaria* guts of Cycle 1 also display a surprising lack of particles >50 µm despite their presence in the water, as shown by FlowCam and epifluorescence microscopy. Similarly, the maximum particle size was only 24.7 µm in the *Thetys vagina* guts from Cycle 1 (Supplementary Table 2). All other salp species regardless of cycle also display this trend, with the exception of *Pegea confoederata* for which the contribution of large particles is still far lower than predicted by the ambient spectra of Cycle 2.

**Figure 4.**
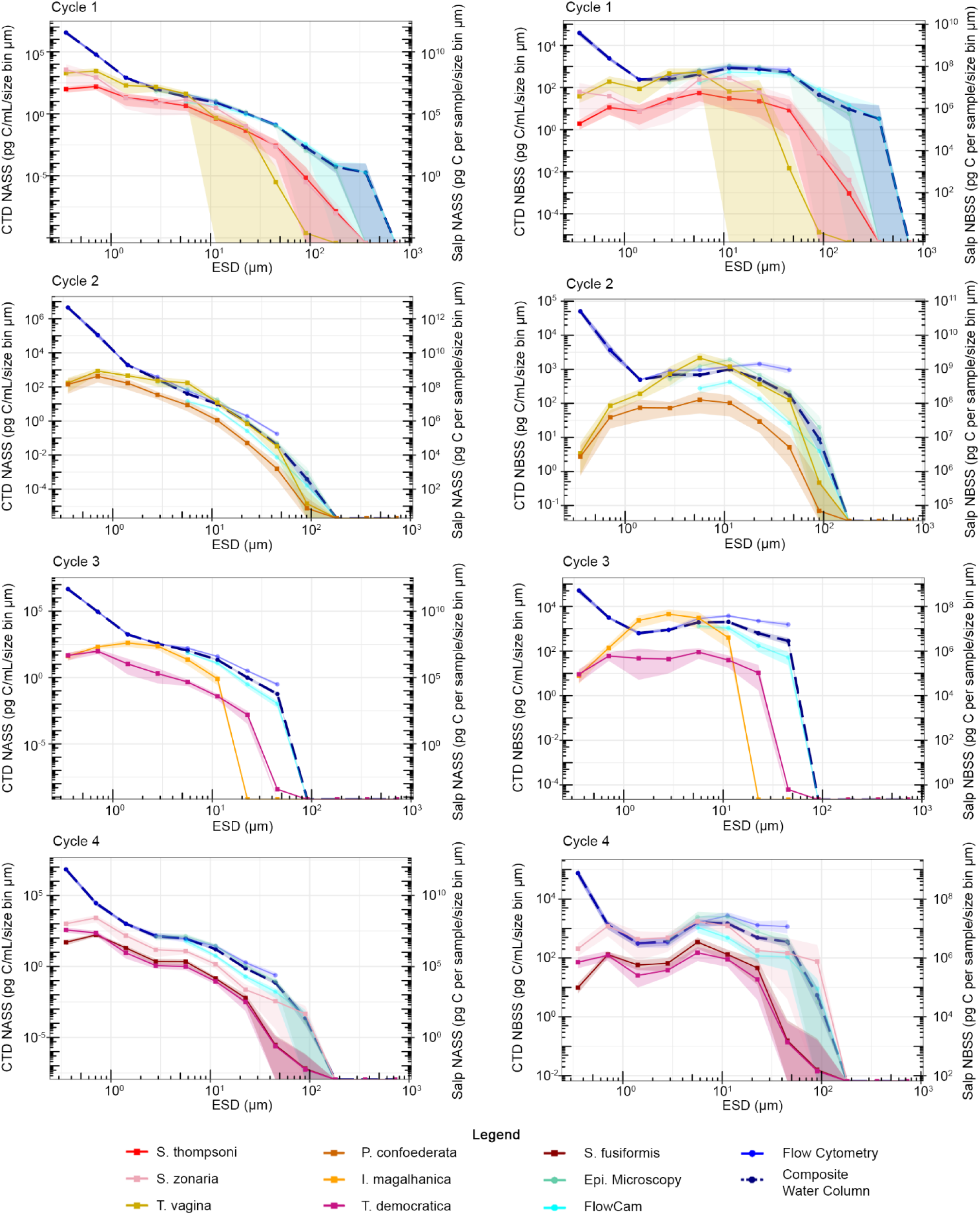
Average normalized abundance (left) and normalized biomass (right) size spectra of salp prey (NASS and NBSS, respectively) as a function of equivalent spherical diameter (ESD) for Cycles 1-4. Warm-colored lines denote salp gut contents while cool-colored lines represent water column measurements from samples collected from the CTD rosette. The thicker, dark blue, dashed line (labeled composite) represents the geometric mean of the water column measurements. Shaded regions represent 95% confidence intervals for each spectrum. Note the difference in units for y-axes and that some water column measurements are not visible due to overlap with the composite spectrum. A version of this figure excluding broken particles can be found in the Supplementary Materials (Supplementary Figure 4).

### 3.2 Feeding distribution parameters

Most salp species obtained the majority of their carbon from particles less than 13 µm (Figure 5; Supplementary Table 2), likely due to the predominance of nanoplankton present in all four cycles coupled with the salps’ apparent difficulty in capturing large particles. Notably, the smallest salp species (*Thalia democratica*) had the second lowest average prey size of 8.4±5.5 µm ranging from 6.2 to 12.0 µm (Figure 6) which may be due to better adaptation for feeding on small particles or simply because the majority of this species was collected from Cycle 3 which had relatively few microplankton (Figure 3). Species from Cycles 1 and 2 (where there were more large particles in the water) such as *Salpa thompsoni* and *Soestia zonaria* typically fed on slightly larger prey, with average prey size ranging from 2.9 to 33.6 µm with a mean of 13.8±8.1 µm for *S. thompsoni* and 10.3 to 23.9 µm with a mean of 14.8±12.4 µm for *S. zonaria*. Within species, the size of the salp did not seem to substantially affect the mean prey size. For example, the mean prey size of *Pegea confoederata* was 10.6±9.0 µm ranging from 3.6 to 13.0 µm for individuals varying in size by almost an order of magnitude. Even between species, no trend with size was observed as the mean prey size of the largest salp *Thetys vagina* is quite comparable at 9.6±9.2 µm with a range of 7.4 to 11.7 µm to that of the smallest species *T. democratica*. Likewise, there does not seem to be any trend with salp life stage as the mean prey size was similar for all species for which we encountered both stages: *S. zonaria* (aggregate=18.2±16.0 µm, n=3; solitary=11.3±8.8 µm, n=3), *P. confoederata* (aggregate=10.8±8.8 um, n=13; solitary=10.2±9.6 µm, n=5), *S. thompsoni* (aggregate=14.2±8.6 µm, n=11; solitary=12.2±6.5 µm, n=3), *T. democratica* (aggregate=7.8±5.3 µm, n=5; solitary=9.2±5.8 µm, n=4). *Ihlea magalhanica* had the smallest mean prey size (4.3±2.5 µm ranging from 3.8 to 4.7 µm) because their guts almost exclusively contained small white spheres and bacteria-like particles. The largest degree of variability in mean prey size was seen in *S. thompsoni*, which had several individuals where more than 50% of their gut biomass was contained in one or two very large particles, such as polycystine radiolarians or large dinoflagellates, substantially inflating the mean prey size. This also occurred with a single *S. zonaria* from Cycle 4, substantially increasing its mean prey size relative to the other samples as well as the average contribution of microplankton to the species’ total gut biomass (Figure 6). *S. thompsoni* was also the most abundant salp species and one of the taxa for which we collected the most samples (n=14) such that the probability of catching a *S. thompsoni* with a large, rare prey particle in its gut was higher.

**Figure 5.**
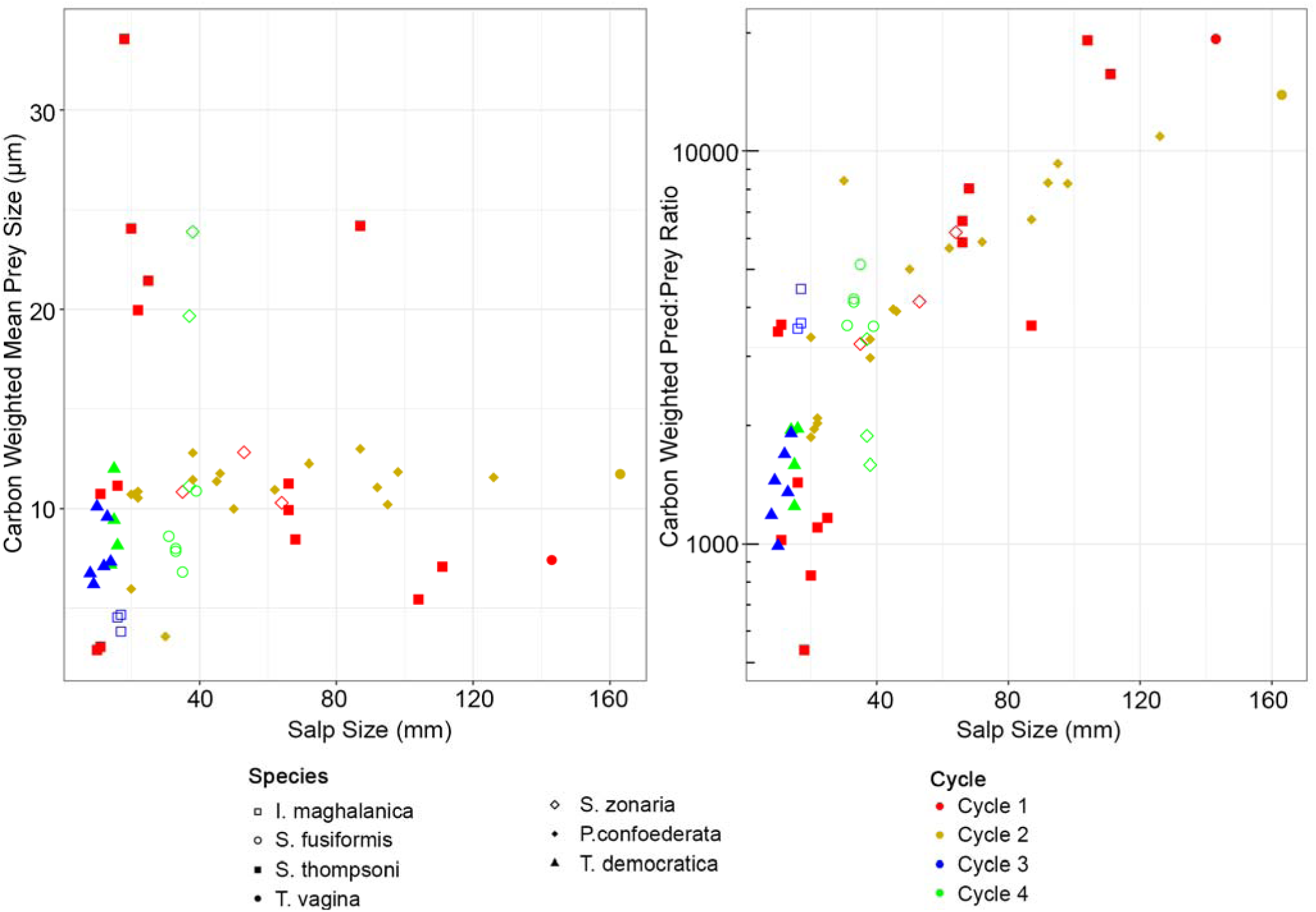
Prey size (left) and predator:prey size ratio (right) at which 50% of gut content biomass is achieved as a function of salp size for each of the 58 salps collected. Shape denotes salp species while color denotes the cycle in which the sample was collected. See Supplementary Table 2 for more information.

**Figure 6.**
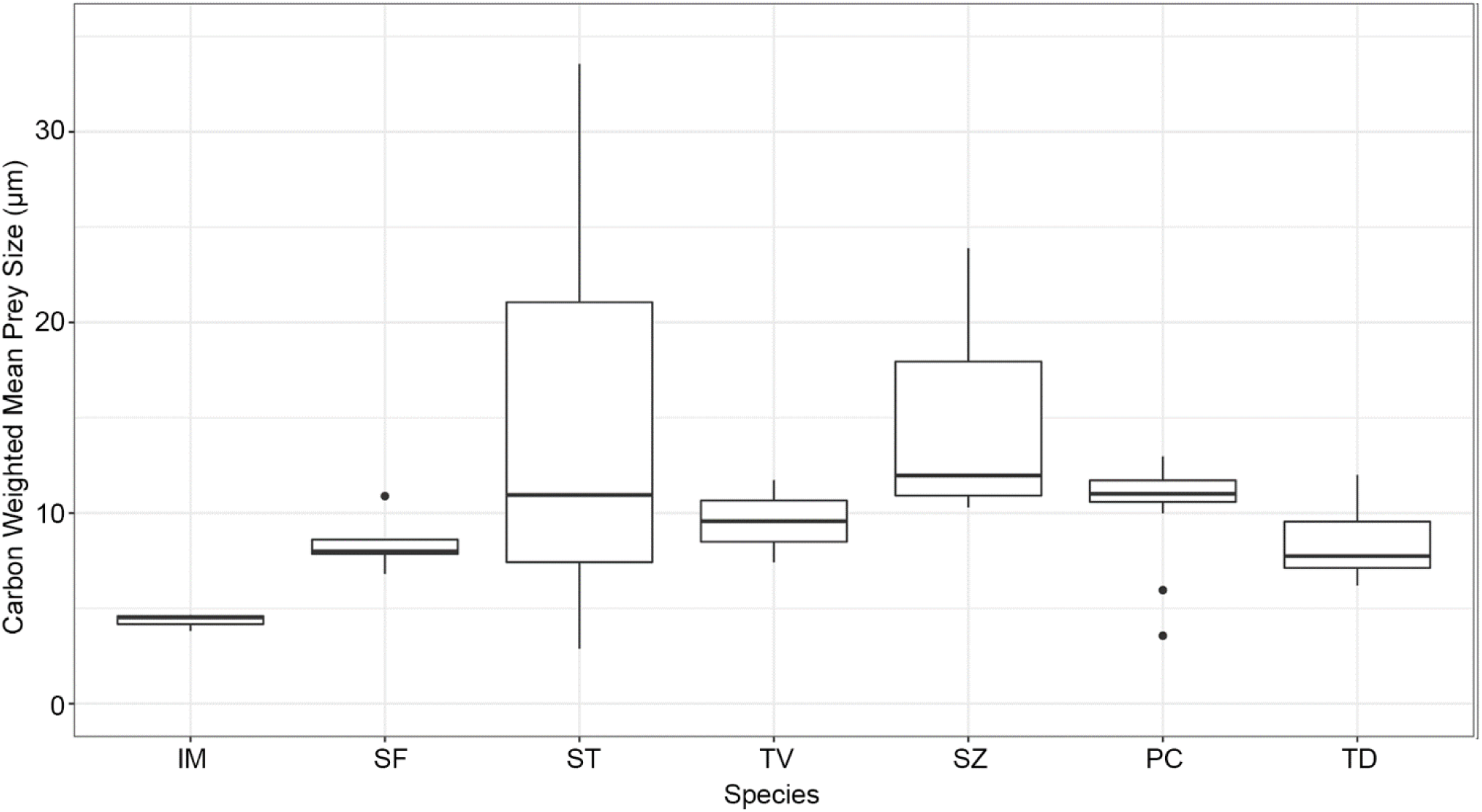
Box and whisker plot of the distribution of carbon-weighted mean prey sizes averaged by salp species. SF = *Salpa fusiformis*, ST = *Salpa thompsoni*, TV = *Thetys vagina*, SZ = *Soestia zonaria*, PC = *Pegea confoederata*, IM = *Ihlea magalhanica*, TD = *Thalia democratica*.

Most salps had predator-prey size ratios (PPSR) between 1,000:1 and 10,000:1, with only the two largest *S. thompsoni* individuals, the two largest *T. vagina*, and the largest *P. cofoederata* feeding at or above 10,000:1 (Figure 5). Averaging across species *T. vagina* had the highest PPSR of 16,365:1, followed by *P. confoederata* (4,475:1), *Salpa fusiformis* (4,093:1), *I. magalhanica* (3,858:1), *S. zonaria* (3,062:1), *S. thompsoni* (2,938:1), and *T. democratica* (1,497:1). When averaging by life stage as well, solitary stage salps always had a higher PPSR than aggregates of the same species, although this may be due to the fact that the solitary stage individuals tended to be larger. There was a clear trend of increasing PPSR with increasing salp size because of the larger variance in salp sizes compared to the relatively similar mean prey sizes. However, a great deal of variation in PPSR was seen amongst individuals of the same species and size. The PPSRs of smaller *S. thompsoni* range by almost an order of magnitude, while *P. confoederata* and *T. democratica* possessed many individuals of approximately the same size yet also show PPSRs that vary by a factor of 2-3 in a given size class, reflecting similar variability in mean prey size. Also of note, PPSR was largely independent of which cycle the salp was collected from; 35-40 mm *S. fusiformis* and *S. zonaria* collected during Cycle 4, which was much farther from the coast and displayed relatively higher concentrations of dinoflagellates were comparable to *S. zonaria* and *P. confoederatea* of the same size from the coastal, diatom rich Cycle 1. We caution that this does not imply that the differences in prey community composition across cycles have no bearing on PPSR, but rather that the greater degree of variability seen in salp size is the primary determinant.

## 4. Discussion

### 4.1 Feeding distribution parameters

The relative abundance of predators with different PPSRs has the potential to substantially alter relationships between trophic level and size, with commensurate impacts on energy transfer to larger taxa of commercial importance and carbon export into the deep ocean (Barnes et al. 2010; Michaels & Silver 1988). In addition to a predator’s mean PPSR, the range of sizes upon which a predator can feed has the potential to structure food webs. Fuchs and Franks (2010) concluded that a dominance of specialized predators which feed on a narrow range of prey sizes (low SD_PPSR_) or are large relative to their prey (high PPSR), such as copepods, would lead to an ecosystem state with lower connectivity between trophic levels, fewer omnivores, more top predators, and greater transfer efficiency to higher trophic levels due to the reduced number of trophic interactions. Conversely, high SD_PPSR_ and/or low PPSR predators like dinoflagellates, raptorial ctenophores, or ciliates led to an ecosystem state with the opposite characteristics.

We carefully quantified the mean PPSR for 7 salp species and the range of sizes that comprised the majority of their carbon-weighted prey, finding Southern Ocean PPSRs that were typically higher than the summary of estimates for salps reported in a meta-analysis by Fuchs and Franks (2010) from other regions. We saw typical values of 1,000:1 to 10,000:1, with all but 3 individual salps >1,000:1 and 62% of our dataset above the maximum reported range of Fuchs and Franks (2010) at 2,236:1 (Figure 7). As salps have usually been regarded as non-selective feeders, some of this difference may be due to the different prey communities present in the regions from which Fuchs and Franks (2010) acquired their data, which include the Mid-Atlantic Bight (Vargas and Madin 2004), Subarctic Pacific (Madin and Purcell 1992), Florida Current, and Gulf of California (Madin 1974). Their data also include several species not observed in our study, such as *Cyclosalpa affinis, C. pinnata, C. bakeri*, and *Weelia cylindrica,* which are quite physiologically distinct from the species investigated here and may also exhibit different feeding characteristics (Harbison and McAlister 1979; Bone et al. 2003). The individuals in the Fuchs and Franks (2010) dataset are also on the small end of our size range, with 3 out of their 5 datasets containing only salps 100 mm (whereas our *T. vagina* were as large as 163 mm). This also likely plays a role in the larger values we observed, because we found little difference in prey size between large and small salps. Considering our Southern Ocean data along with the data from other regions described above, salp PPSRs range over two orders of magnitude, which is comparable to the difference between pallium-feeding dinoflagellates and suspension-feeding copepods. The SD_PPSR_ for our study ranged from 0.15 to 0.50, while Fuchs and Franks (2010) estimated similar values ranging from 0.22 to 0.50. Thus, the range of prey sizes salps feed on can be broader than most other planktonic predators or as narrow as the protists they compete with, again largely varying with the specific conditions of a given study region. Attempts to characterize salp predator:prey interactions in models using a single set of generalized parameterizations would fail to capture this regional variability and may lead to significant error when applied to large, heterogeneous areas such as the Southern Ocean. Even for global modelling attempts, it is clear that ascribing a single PPSR of 10,000:1 will lead to misleading results. Models should endeavor to use appropriate PPSRs depending upon the prey field and salp population in the region studied.

**Figure 7.**
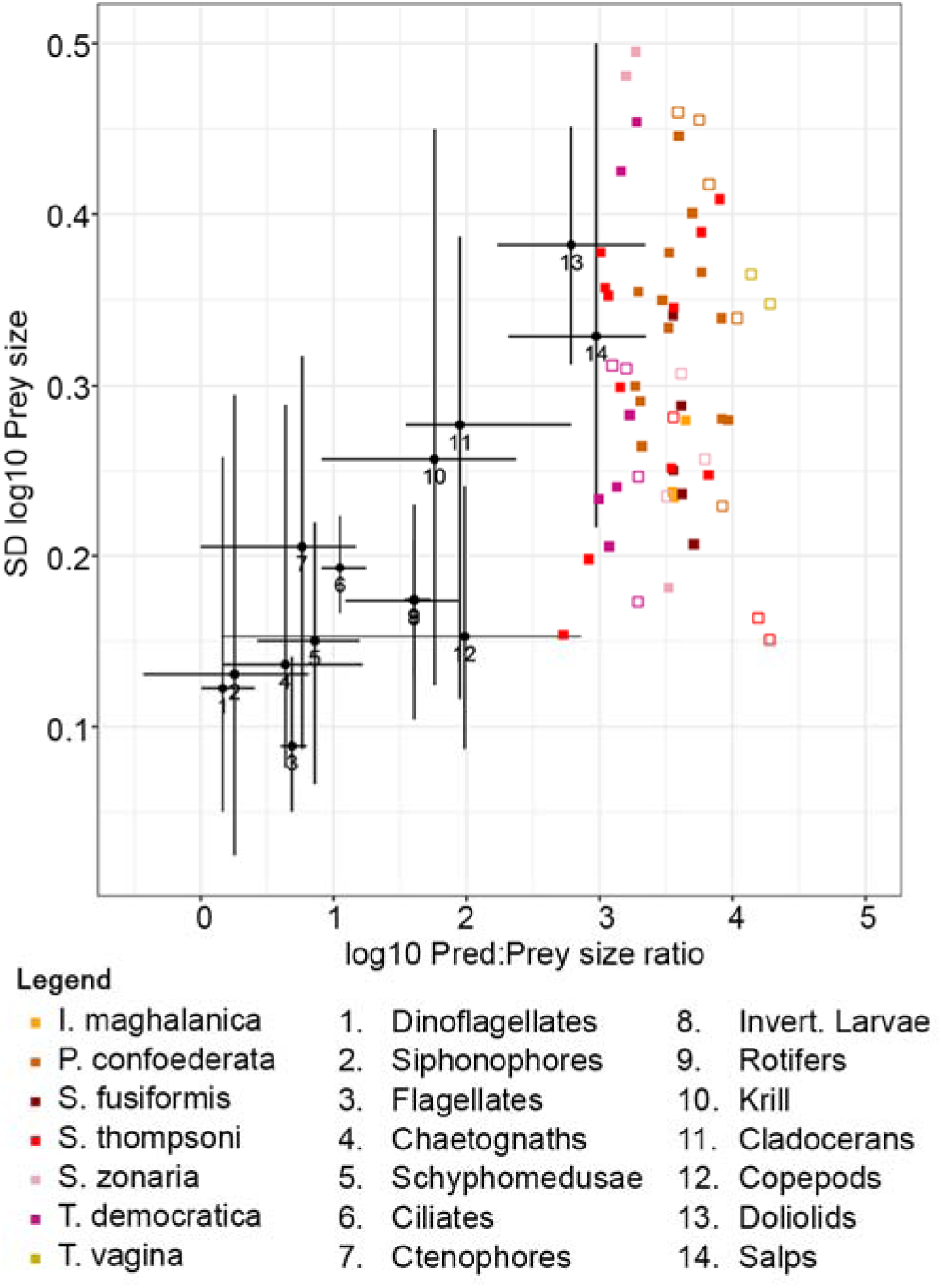
Predator:prey size ratios and standard deviation of prey size for a variety of planktonic predators from Fuchs and Frank (2010; black circles) with bars representing the range. Squares represent individual salp data from this study, with color representing salp species. Filled symbols denote aggregates and empty symbols denote solitaries. Note that values from this study are carbon-weighted whereas those of Fuchs and Franks (2010) are not.

While our estimates of SD_PPSR_ are fairly comparable to those of Fuchs and Franks (2010), 14% of our dataset is below their minimum value of 0.22. This is probably due in part to the dominance of ∼10 µm prey particles that were ubiquitous across nearly all salp’s guts sampled in our study (Figure 5). While many of these were clearly small dinoflagellates or centric diatoms, the most common particulate items found in salp guts were ∼2-7 µm smooth white spheres (Supplementary Fig. 3H). These spheres made up 30% of all identified particles and were present even in samples where little to no other recognizable taxa were found. Unknown particles matching this description were also described by both Madin and Purcell (1992) and Ahmad-Ishak et al. (personal communication) and can be seen in the SEM images taken by Caron et al. (1989) although they were not discussed. Due to their size and generally spherical shape, we assume that the majority of these particles were nanophytoplankton such as prasinophytes, prymnesiophytes, or pelagophytes for which characteristic features such as flagella had been digested or broken off as these groups had a high abundance and contribution to phytoplankton community biomass (Décima et al. 2023). It is, however, likely that this morphological categorization includes particles of various origins. For example, some displayed significant silicon signals under electron diffusion spectroscopy and therefore may have instead been the resting stage cysts of diatoms. Alternatively, some may have been debris or tissue associated with the preparation of the salp itself rather than ingested prey particles. How these particles are treated has a strong impact on how some of our results should be interpreted. For example, the NBSS of *I. magalhanica* showed a strong peak at 2-8 µm due to the exceptionally high numbers of white spheres observed in the guts of all 3 individuals imaged (Figure 4). When these are treated as nanoplankton, as we believe is most likely, our data suggests feeding on particles of this size class is even more important for this species of salp relative to the others investigated here. If white spheres were not treated as particles and instead excluded from our analysis, the resulting spectra for *I. magalhanica* would be much the same as that of *T. democratica* from the same cycle and would instead suggest their feeding dynamics are similar (Supplementary Figure 5). Both salp spectra, however, would then be significantly lacking in particles of this size relative to the ambient spectra. This is also true of the other salp spectra across all cycles and species, with even more noticeable declines in nanoplankton for the salps of Cycles 2 and 4. We interpret this as evidence that the majority of these white spheres indeed originated in the water column but caution that future work, especially if seeking to automate image processing, should take special care with particles in this size class.

### 4.2 Retention efficiency

For most salps, we find near 100% retention for 1-16 µm particles (Figure 8), but much lower retention for submicron particles, which agrees with many previous studies (Harbison and Gilmer 1976; Kremer and Madin 1982; Caron et al. 1989; Sutherland et al. 2010; Nishikawa and Tsuda 2021; Stukel et al. 2021). With respect to the larger prey size classes, however, the most striking result of our calculations is the rapid decline in retention of 16-64 µm particles for many salps. While our methods were optimized for finding the lower threshold of retention, making it difficult to pinpoint a single particle size for the drop-off, it is clear that this resulted in surprisingly low contributions of microplankton to the total salp gut biomass compared to what was found in the water column.

**Figure 8.**
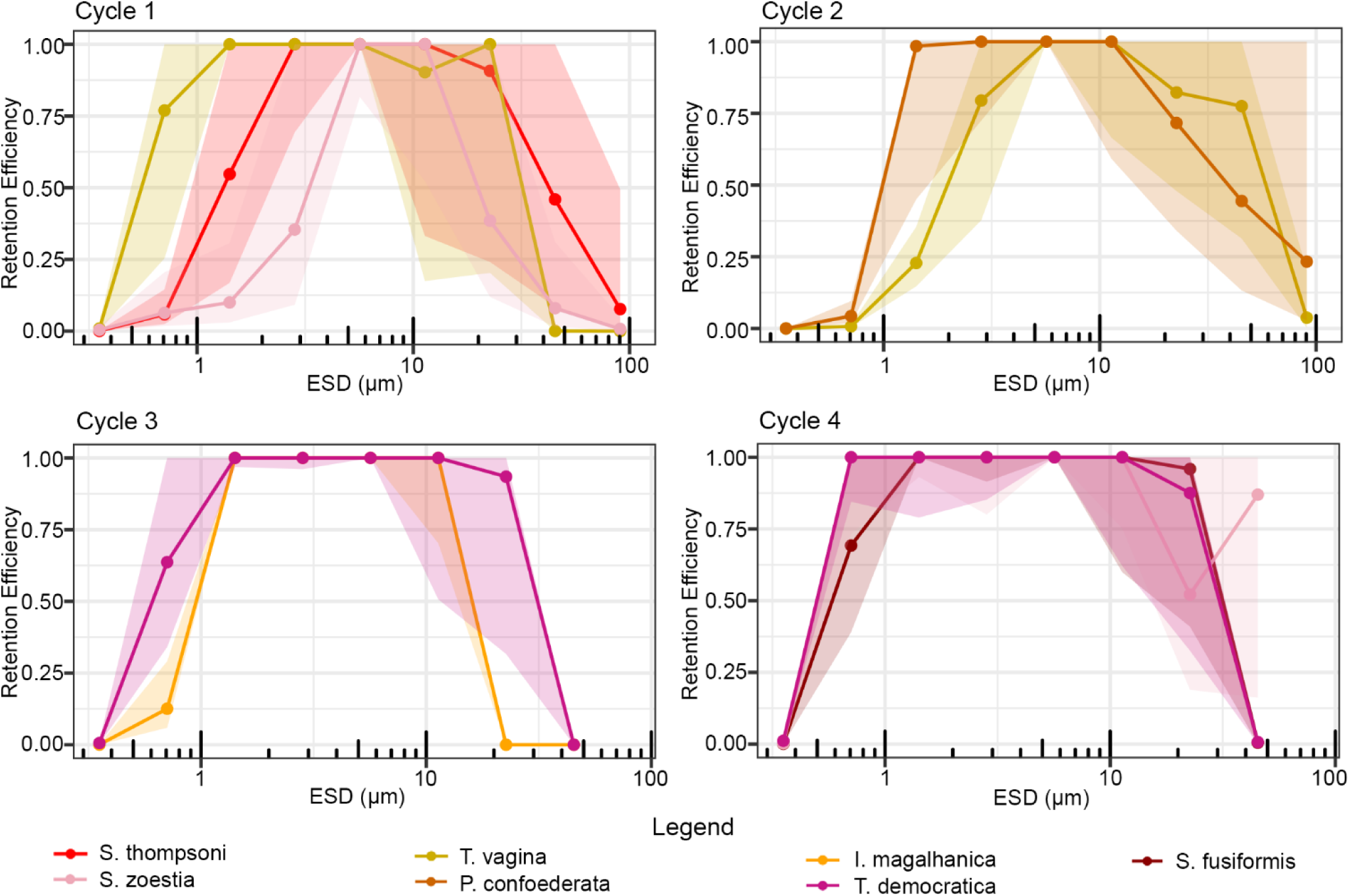
Average retention efficiency as a function of prey equivalent spherical diameter (ESD) for each salp species, organized by cycle, assuming filtration rate is equivalent to the clearance rate on 8-32 µm cells. Shaded areas represent 95% confidence intervals. A version of this figure excluding broken particles can be found in the Supplementary Materials (Supplementary Figure 6).

Typically, all particles above the width of a salp’s mucous feeding mesh are considered to be retained with 100% efficiency. However, previous studies frequently used particles of only up to ∼10 µm in size, so little data existed for larger particles and their relative absence from salp diets may have been easily missed. Alternatively, several factors could have led to an underestimate of RE for these larger size classes in the present study. Solitary *Nitzschia* and *Pseudonitzschia*-like diatoms made up the majority of the pennates identified in the salp guts, which stands in contrast to the many-celled chains of these genera observed in the FlowCam. It is possible that larger, fragile organisms like these chains of pennate diatoms were separated into solitary cells and/or further fragmented as the salp’s filter was passed through the esophagus and into the stomach which would bias the prey size spectra small in comparison to what was observed in the field. It should be noted that this fragmentation may also have occurred as a result of our preparations in the lab, particularly vortex mixing and vacuum filtration. Indeed, deck-board prey disappearance experiments with *S. thompsoni* during our cruise, which would not suffer this bias, did not show a consistent decrease in RE for particles up to 30 µm ESD (Stukel et al. 2021). However, this trend remains even after correction for particle and chain breakage, which suggests that these large particles were truly lacking in the salps’ guts. The incubation experiments of Vargas and Madin (2004) also found 30-40% lower retention efficiencies for 60 µm diatoms in Mediterranean *Salpa cylindrica* and *Cyclosalpa affinis*. One potential explanation for this revolves around “clogging” whereby a bolus forms in the salp’s feeding filter and prevents it from being ingested. What little work that has been done investigating this phenomenon has focused on the role of particle abundance (Harbison and Gilmer 1976; Harbison et al. 1986), but it has also been noted that particle type may be important. Vargas and Madin (2004) noted that clogging could explain the lack of large diatoms in their salps, a trend we also observe. Our results may support the existence of a size threshold for large particles at which the likelihood of bolus formation and subsequent expulsion of the feeding filter further decreases their already low relative abundance in the guts. To this end we noted a complete absence across all 58 salps of large or spiny centric diatoms such as *Chaetoceros* sp. (both solitary cells and chains) as well as those which appeared to be either *Proboscia* sp. or *Rhizosolenia* sp., all of which were common contributors to the >60 µm size classes in the FlowCam. Similar discrepancies were observed by Ahmad-Ishak et al. (2017) who noted that while both *Chaetoceros* and *Proboscia* made up large biovolumes in their study region according to light microscopy, only *Proboscia alata* was found in the guts of *S. fusiformis* and *T. democratica* via SEM. Unfortunately, our understanding of the conditions under which boluses form is still too poor to allow for more than speculation here and further investigation is warranted.

Systematic differences in mesh size based on species is one potential explanation for the markedly higher submicron retention efficiencies of some of the salps investigated here (Figure 8). Only a handful of studies have attempted to quantify the mesh spacing of the mucous feeding filter in just a few species of salps, yet the variability in mesh spacing reported supports this possibility (e.g. 1.9 x 0.2 µm in *S. fusiformis*, Silver and Bruland 1981; 0.7 x 4.0 µm in *P. confoederata*, Bone et al. 1991; 1.3 x 1.3 µm in *S. fusiformis*, Bone et al. 2000; 0.9 x 0.9 µm in *T. democratica*, Bone et al. 2003). These few studies are likely the result of the difficulty of preparing these fragile filters (typically from singular animals) and thus it is difficult to determine which factors, including species, ultimately impact mesh spacing. For example, Bone et al (2003) compared direct observations of filter meshes across multiple species, studies, and preparatory methods and found that dimensions are likely subject to a variety of factors such as fiber thickness, elasticity, strength of the inhalant current, and even location within the salp. Thus further work will be necessary to determine which factors are the most important for understanding salp mesh sizes and their implications for prey selection.

The trends regarding retention efficiency with salp size are thankfully more straightforward. Higher retention of small particles in smaller salps has been reported by several investigators (Harbison and McCalister 1979; Kremer and Madin, 1992, Stukel et al. 2021) potentially due to isometric scaling of the mesh width to organism size (Sutherland et al. 2010). Our results indicate that *T. democratica*, for example, have the highest RE for submicron particles when averaged over all samples (81.9% for 0.5-1 µm particles, Figure 8) which is in agreement with field reports of efficient retention of bacteria in this species (Mullin 1983; Vargas and Madin 2004). They are also the smallest species we investigated, with an average size of only 12.6 mm. Similarly, *S. fusiformis* REs for submicron particles are significantly higher than the congeneric *S. thompsoni* which struggled to retain anything smaller than 1 µm. We postulate that this may be due less to difference in species and more to the lower mean size of *S. fusiformis* (34.7 mm) captured compared to that of our *S. thompsoni* (50.3 mm). When instead averaging *S. thompsoni* retention across different salp size classes, we again find smaller individuals display higher REs for submicron particles (Figure 9). These observations match reasonably well to those predicted by the equation for *S. thompsoni* size specific clearance rate given in Stukel et al. (2021), which describes mesh diameter as an allometrically-scaling function of salp size. Note that these results do not contradict our earlier observations of mean prey size being relatively independent of salp size. As mean prey size is a function of both retention efficiency and the prey field available, the impact of the differences in RE for small particles we find here on mean prey size, and thus PPSR, will be proportional to that size classes contribution to the ambient carbon pool. Submicron particles never comprised >27.6% (and usually substantially less) of the total carbon in the water column, so it’s not surprising that the mean prey size and PPSR of any given salp was not strongly impacted by its ability to retain these particles.

**Figure 9.**
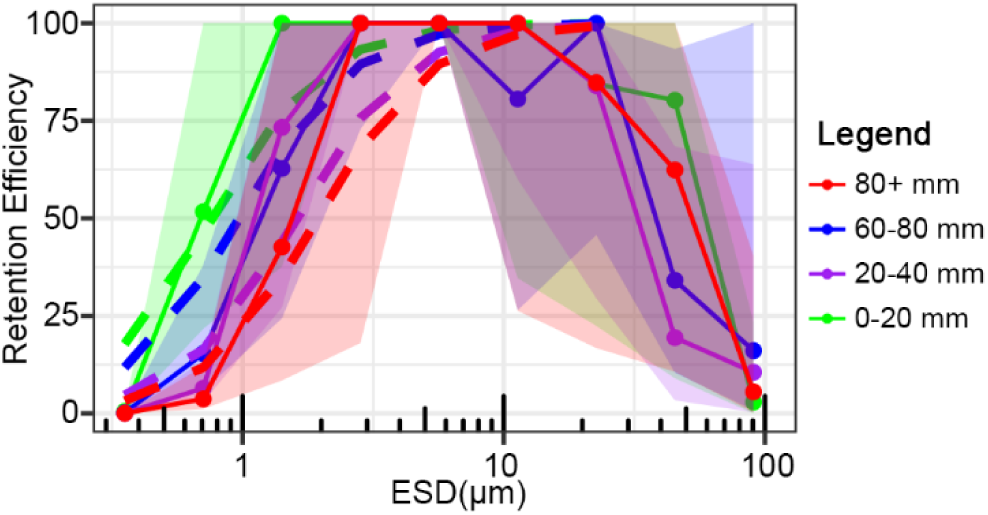
Retention Efficiencies (RE) for different sizes of *Salpa thompsoni* assuming a filtration rate equal to the clearance rate for 8-32 um particles. Solid lines represent REs averaged by salp size class, dashed lines represent REs calculated using the size resolved clearance rate equation from Stukel et al. (2021) for *S. thompsoni*, and shaded intervals represent 95% confidence intervals.

### 4.3 Ecological implications

In this study, we quantified the size spectra of prey items within 7 different species of salps of both life stages and a range of sizes in order to compare them against that of the ambient water column in each of 4 distinct water masses within the Chatham Rise. Our results suggest that these gut size spectra are largely similar to the ambient in most cases, especially within the nanoplanktonic size range (Figure 4), with several notable exceptions. In particular, the salps of Cycle 1 exhibited a far greater contribution of nanoplankton than microplankton to their gut biomass regardless of salp size, species, or life stage than would be expected for a non-selective filter-feeder based on the dominance of microplankton in the prey field surrounding them (Figure 3). This larger contribution of nanoplanktonic prey was also true of the salps from Cycles 3 and 4, although the contribution of microplankton in the prey field during these cycles was lower than in Cycle 1. While these observations may support growing evidence in favor of selective feeding in salps (von Harbou et al. 2011; Metfies et al. 2014; Dadon-Pilosof et al. 2019; Pauli et al. 2021; Thompson et al. 2023), similar differences in prey contributions for doliolids have been linked to small-scale heterogeneity in the prey community due to micro patches and thin layers (Takahashi et al. 2015; Walters et al. 2019; Greer et al. 2020; Frischer et al. 2021). Our vertically-integrated ambient prey abundances fundamentally average over these fine-scale features. Since the net deployments used to collect salp samples integrated over up to 200 meters, it is possible the prey field fed upon and therefore represented in their guts differs to some degree from the average field represented by the water column data. Picoplankton also made up far less of the salp gut biomass than that of the water column in Cycles 1, 2, and 4, perhaps due to the salps’ difficulty in retaining submicron particles as well as the low contribution of picoplankton to total biomass. In contrast, picoplankton biomass in salp guts from Cycle 3 was about equal to that of the water column. While this is likely due to the predominance of nanoflagellates in the guts of *Ihlea magalhanica*, for *Thalia democratica* it suggests more efficient retention of submicron particles than other salp species. Mean prey sizes, while highly variable, followed similar patterns with larger average prey sizes for salps from the subantarctic Cycles 1 and 2 and smaller prey sizes for Cycles 3 and 4 (Figure 5).

Variation in how differently sized particles are consumed by salps is of great ecological importance as it determines how salp feeding structures the plankton community around them, but the contribution of different size classes to the total biomass ingested by the salp is also relevant from the perspective of salp energetics. As our results have shown, the latter is not exclusively predicated on the former and, as in all filter feeders, also depends on the prey size structure available to feed on. For example, even if inefficient submicron feeding results in larger numbers of bacteria being removed from the water than traditionally believed (Sutherland et al. 2010; Dadon-Pilosof et al. 2019) and smaller salps do indeed display more efficient retention for submicron particles (Harbison and McCalister 1979; Kremer and Madin, 1992, Stukel et al. 2021), the lower total carbon content per cell attributable to small particles compared to those that are larger and rarer but more efficiently retained and carbon-rich means these submicron particles may still be relatively inconsequential to a salp’s diet. Indeed, we found large numbers of bacteria-like submicron particles in the salp’s guts, however <1 µm particles made up only 2.5±3.4% of the average salps gut content. Consequently, the contributions by picoplankton to salp gut content were lower than their contribution to the water column in 93% of the individuals sampled (Figure 3). Most salps instead showed carbon-weighted mean prey sizes of 6-13 µm (Figure 5), which agrees with Stukel et al.’s (2021) results for *S. thompsoni* that found carbon-weighted median prey sizes of 8-9 µm across all cycles.

Likewise, a variety of reports suggest that small diatoms and dinoflagellates make up the bulk of the diet for many different species of salps (Vargas and Madin 2004; Tanimura et al. 2008; Ahmad-Ishak et al. 2017), regardless of sampling location or season (von Harbou et al. 2011), implying this ∼10 µm size class may represent a sweet spot between numerical abundance in the water column and carbon content. While diatoms and thecate dinoflagellates contain hard frustules or theca that are often resistant to digestion and may therefore be expected to contribute disproportionately to gut contents, molecular and fatty acid composition analyses of *S. thompsoni* and *I. racovitzai* in the Southern Ocean also found higher abundances of dinoflagellates and diatoms in the guts (von Harbou et al. 2011; Metfies et al. 2014; Pauli et al. 2021). This contrasts with the findings of Sutherland et al. (2010), who reported submicron particles alone could make up more than 100% of the carbon requirement for *P. confoederata*. However, this conclusion depended on their assumption that only the outer 0.1 µm of ingested cells were digested. When instead assuming complete digestion of all particles, as we do here, Sutherland et al. (2010) also found the majority of ingested carbon came from 1-10 µm particles such as nanoflagellates and small diatoms. The true degree of digestion will vary based on many characteristics of a given particle and the salp’s gut turnover time. Until such time as when we can better predict the nutritional values for different plankton types, our results suggest that the true mean prey size of salps in the Southern Ocean is close to 10 µm.

## 5. Conclusion

Overall, our results indicate that within the 1-16 µm prey size range, salp diets for the seven species investigated here largely reflect the size composition of plankton in the surrounding water regardless of species, size, or life stage. Feeding on submicron particles, however, appears to be dependent on both salp species and/or size. *T. democratica*, perhaps due to its smaller average size, was able to consistently retain even submicron particles efficiently, whereas most of the larger species such as *T. vagina*, *S. thompsoni*, and *P. confoederata* showed reduced retention efficiencies below 1 µm. Size resolved retention efficiencies for *S. thompsoni* also showed this trend. Retention for particles >1 µm was generally high, with a decrease in efficiency for >16 µm particles across all salp species studied, possibly as a result of difficulty ingesting larger particles. Coupled to their prevalence in the water column, this caused nanoplankton to comprise the majority of the carbon in salp guts across all samples. This led to predator:prey size ratios ranging from 536:1 for small *S. thompsoni* to 19,285:1 for large *T. vagina* with most falling between 1,000:1 and 10,000:1. Rather than being due to systematic differences in filtration physiology, however, our results indicate this order of magnitude variability is primarily due to the large range of sizes over which salps can occur as well as the spectra of the ambient prey field. In other words, while the ability of salps to feed on small particles is variable and ecologically important, this variation is less important with respect to salp energetics. Future work to investigate prey size spectra across a broader range of salp sizes and species is necessary to disentangle the potentially confounding impacts of these factors.

## 7. Acknowledgements

We would like to thank the captain and crew of the R/V Tangaroa for help in the deployment of equipment as well as our many collaborators in the SalpPOOP project including Scott Nodder, Sadie Mills, Florian Lüskow, Morgan Meyers, Sarah Searson, Lana Young, Siobhan O’Connor, Karl Safi, Adriana Lopes dos Santos, and Fenella Deans.

## 9. Data Availability

All data is accessible on the Biological and Chemical Oceanography Data Management Office website (https://www.bco-dmo.org/project/754878).

## 11. Ethical Approval

All applicable international, national, and/or institutional guidelines for sampling and experimental use of organisms for the study have been followed and all necessary approvals have been obtained.

## 12. Author Contributions

CKF and MRS wrote the manuscript and conducted the data analysis. MD was responsible for cruise planning and locating salps. KS conducted FlowCam and flow cytometry sampling. NY conducted epifluorescence microscopy sampling and image analysis. MD and CKF were responsible for net deployments and sample collection. CKF was responsible for SEM and FlowCam image analysis. All authors contributed to editing the manuscript.

