## Supplementary Materials for "Prey size spectra and predator to prey size ratios of southern ocean salps"

**2. Methods**

**2.3.1 Taxon-specific Image Processing**

Four categories of particles imaged by the FlowCam proved problematic due to poorer outline quality and had to be treated specially. Many pennate diatoms, for example, were largely transparent such that VisualSpreadsheet could only detect the particle’s edge from the background in the image (Supplementary Figure 1 A1 & A2). This resulted in adequate estimates of length, but extreme underestimates of width. To remedy this, we manually measured the caliper width of every such pennate diatom in the first CTD cast of each of the four cycles to derive the average pennate width for that cast (n = 104, 37, 6, and 20 for Cycles 1-4 respectively), and applied this average retroactively to all poorly detected pennates within that cycle. Equivalent spherical diameter (ESD) and biovolume (BV) for the resulting prolate spheroid were calculated using Equations 1 and 2 (main text). A similar problem was encountered with some semi-transparent dinoflagellates (Supplementary Figure 1 D1 & D2). For these we re-calculated the width using the recorded length and mean aspect ratio for well-detected dinoflagellates in the first CTD cast of each cycle and again used Equations 1 and 2 for ESD and BV. The spiny protrusions of *Chaetoceros* (Supplementary Figure 1 B1 & B2), however, required we take a different approach to avoid fitting a spheroid that greatly overestimated BV. Instead, we recalculated the length and width of these particles using the recorded aspect ratio and area (n.b., holes in the cell outline were rarely a problem for this class). Finally, the helical colonies of *Asterionelopsis* (Supplementary Figure 1 C1 & C2) proved exceptionally challenging. These exhibited all of the problems mentioned above such that the outlines provided were often barely representative of the colony and were not informative. Each *Asterionelopsis* colony was therefore saved as an individual image file (tif format), and representative single cells within a colony that were parallel to the camera’s field of view were manually analyzed using ImageJ (v. 1.52a). This procedure was repeated for all colonies in the first cast of each cycle and the mean width and length taken to produce an “average” *Asterionelopsis* cell. We then manually counted the number of individual cells present in each image and applied them to the averaged cell sizes. While this should result in a relatively accurate summed BV for this phytoplankton class, we caution that the abundance, NBSS, and NASS data will no longer be representative of whole colonies and will therefore be minimum estimates.

**2.3.2 Carbon Conversions**

Since we could not differentiate between different types of prokaryotes within salp guts, we instead chose to estimate an average carbon content for all bacteria-like particles based on the ratio of the three most dominant bacterioplankton groups, namely *Prochlorococcus sp.*, *Synechococcus sp.*, and heterotrophic bacteria, to each other. This ratio was calculated separately for each of the four cycles using abundance estimates from flow cytometry. For a given cycle we then calculate the average carbon content $C_{BLP}$ in pg C μm^-3^ as

4)	$C_{BLP}= {(F}_{Pro}*C_{Pro})+{(F}_{Syn}*C_{Syn})+{(F}_{HBact}*C_{HBact})$

where $F_{Pro}$ is the fraction of the total bacterial community made up by *Prochlorococcus*, $C_{Pro}$ (0.235 pg C μm^-3^; Garrison et al. 2000) is the average carbon content of *Prochlorococcus* from published allometric relationships, and so on for the other two groups ($C_{Syn}$ = 0.235 pg C μm^-3^, $C_{HBact}$ = 0.380 pg C μm^-3^).

**2.4.1 95% Confidence Interval Calculations**

As stated in the main text, all 95% confidence intervals (CIs) presented in figures were calculated using Markov Chain Monte Carlo random resampling of the original dataset to simulate uncertainty in our collection and/or analysis methods. For example, uncertainty in NASS and NBSS estimates for cycle averaged salp spectra were found by randomly selecting individuals from the original pool of salp samples collected in a given cycle, randomly selecting 20 of the SEM images taken at each of the three levels of magnification, and repeating the process for each of the resampled salps to generate a new particle dataset. The NASS and NBSS for this new dataset were calculated and the entire process was repeated a total of 10,000 times or until the results converged. The 2.5^th^ percentile and 97.5^th^ percentile of the 10,000 NASS/NBSS estimates represent our lower and upper confidence intervals respectively. A similar process was conducted for the FlowCam, epifluorescence microscopy, and flow cytometry with the exception that the random resampling was done at the level of choosing CTD casts from a given cycle and then either resampling the 20 images taken as with the salps in the case of epifluorescence microscopy or resampling all of the particles themselves in the case of FlowCam.

For the water column composite spectra, confidence intervals were calculated differently due to the complexity associated with using the original component spectra. Within a given size range, we instead chose to randomly generate new lognormally distributed datasets of 100,000 values with the same mean and confidence intervals as each of the original component spectra (FlowCam, flow cytometry, and/or epifluoresence microscopy) relevant at that size range. We then randomly sampled one value from each of these datasets and took the geometric mean, repeating this process 10,000 times and using the results to determine upper and lower confidence limits as above. Confidence intervals were similarly calculated for the retention efficiencies of each salp species per cycle by generating lognormal datasets for each salp species’ spectra and that of the water column composite spectra per size bin.

**2.4.2 Broken and Disrupted Particle Corrections**

Due to the lack of chained diatoms and high proportion of broken particles in the salp guts, we chose to apply a correction to each of these particles to estimate what their true size likely was at the time of ingestion. This process was conducted as part of the aforementioned Monte Carlo random resampling scheme to fold additional introduced uncertainty in these assignments into our estimates. As a first step, the abundance and biovolume of all intact and broken solitary and chain-forming diatoms observed in the salp guts were reassigned to a size class in proportion to the size distribution observed in the water columns, as observed via FlowCam. For example, if a 10 µm chain-forming pennate diatom was observed in the guts of a salp from Cycle 1, and the relative proportion of intact solitary and chained pennate diatoms in the water column of Cycle 1 was 10, 20, and 70% 5, 10, and 50 µm particles respectively, that diatom would then be assigned to one of those three size bins with likelihood weighted by this distribution. To avoid biasing the total biomass contributions by inflating the volume of broken particles, the originally calculated biovolume was then applied to the assigned size bin rather than that of the particle’s observed size bin. To correct the abundance, we assumed that the diatom represents a fraction of a particle. We estimate this fraction by correlating the observed biovolume to that of the water column average biovolume of all diatoms in the new size class to which the diatom in the gut has been moved. Thus, a broken 10 µm pennate diatom with biovolume 105 µm^3^ would be counted as 4% of a single particle if sorted into the 50 µm size bin with an average biovolume of 2617 µm^3^. This process was repeated for all other categories of broken particles as well, deriving the weighted distribution whenever possible from FlowCam data or otherwise from non-broken particles of the same class within salp guts of the same cycle.

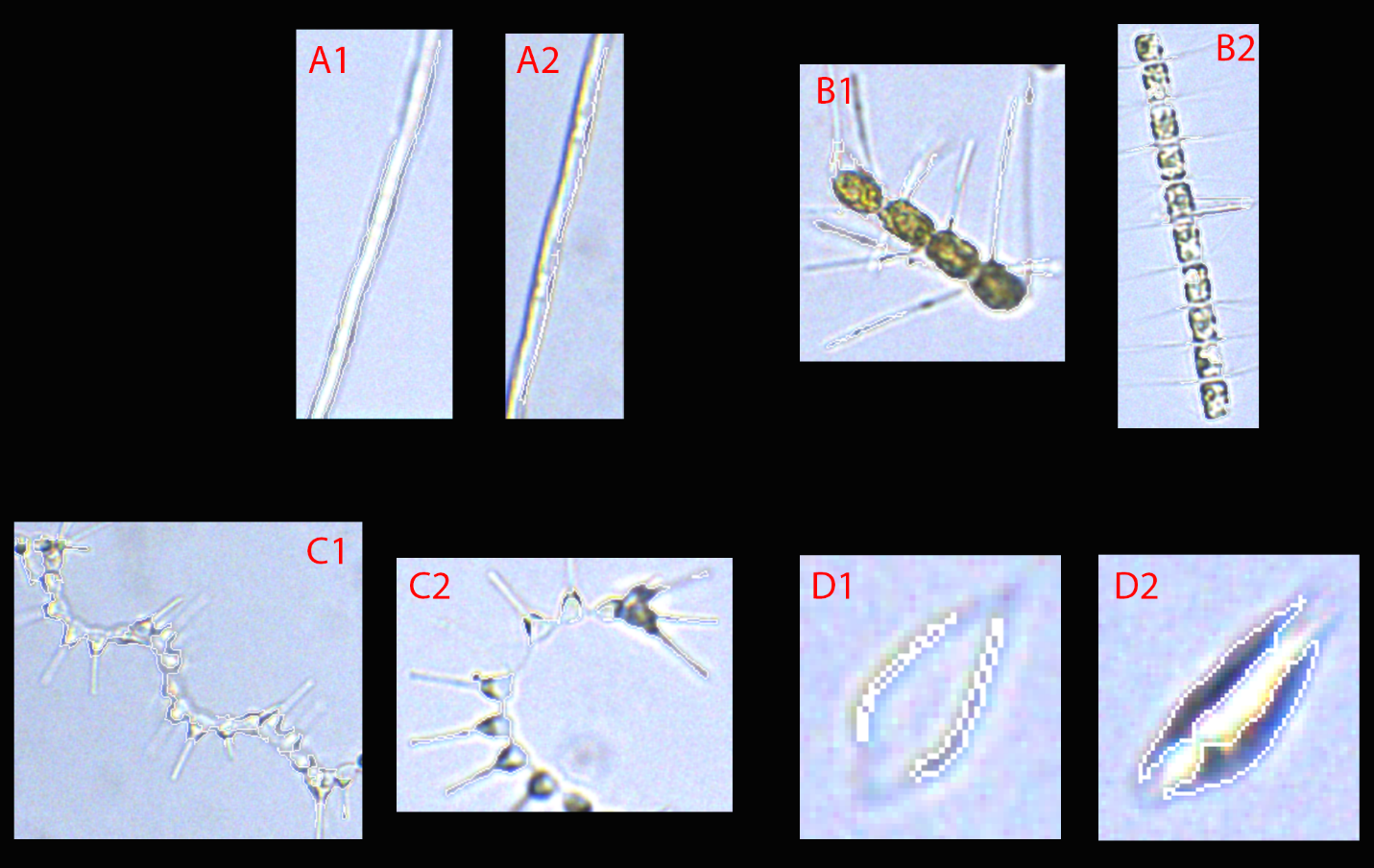

Supplementary Figure 1. Representative images of taxa which were poorly detected by FlowCam. A1 & A2 are pennate diatoms, B1 & B2 are *Chaetoceros*, C1 & C2 are *Asterionellopsis*, and D1 & D2 are transparent dinoflagellates. White outlines in each image represent the region identified by Visualspreadsheet as the particle. Note that images are note to scale.

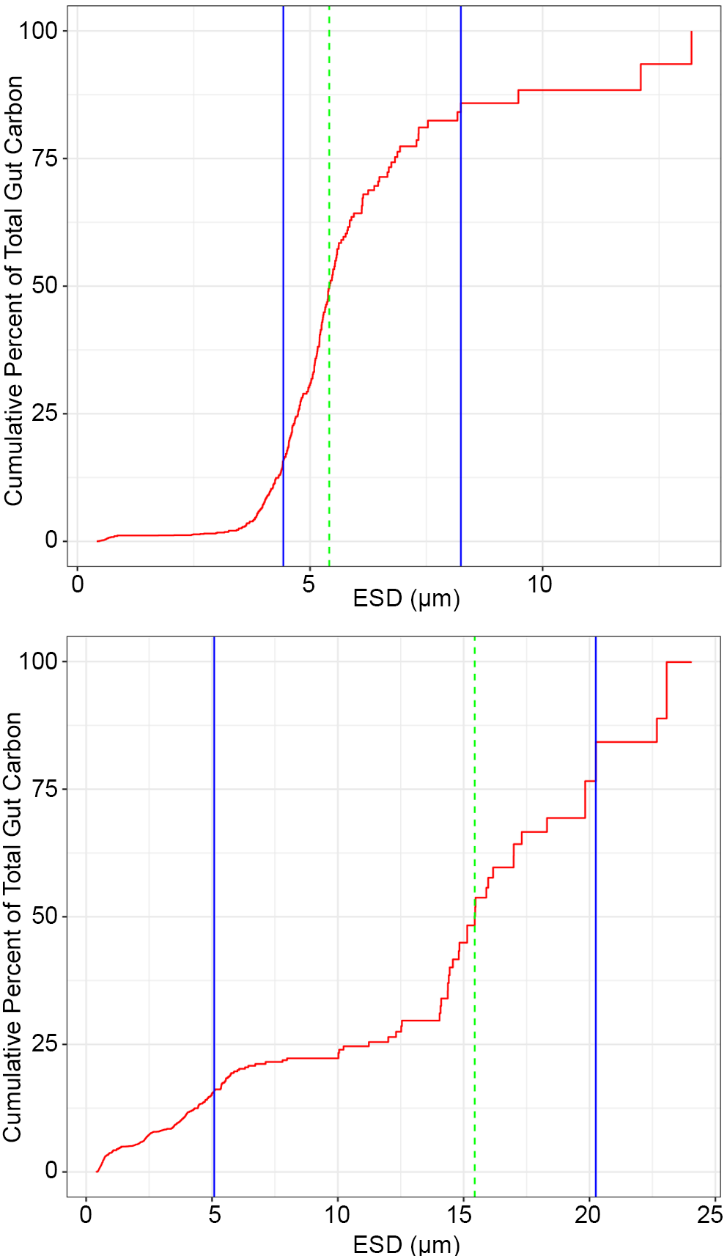

Supplementary Figure 2. Cumulative percent of the total carbon made up by non-broken particles of a given size in the guts of two *S. thompsoni*. Blue solid lines represent the prey sizes at which 15.9% and 84.1% of the total gut carbon has been reached. The log transformed difference of these two values divided by two gives the SD_PPSR_. The dashed green line represents the carbon-weighted mean prey size, or the size of the particle at which 50% of the total gut carbon is achieved. The top salp has a smaller mean prey size with fewer large particles in the gut, resulting in a sharper increase in total gut carbon such that the SD_PPSR_ (or the difference of the two blue lines after log transformation divided by 2) will be lower than the SD_PPSR_ of the bottom salp for which the opposite is true.

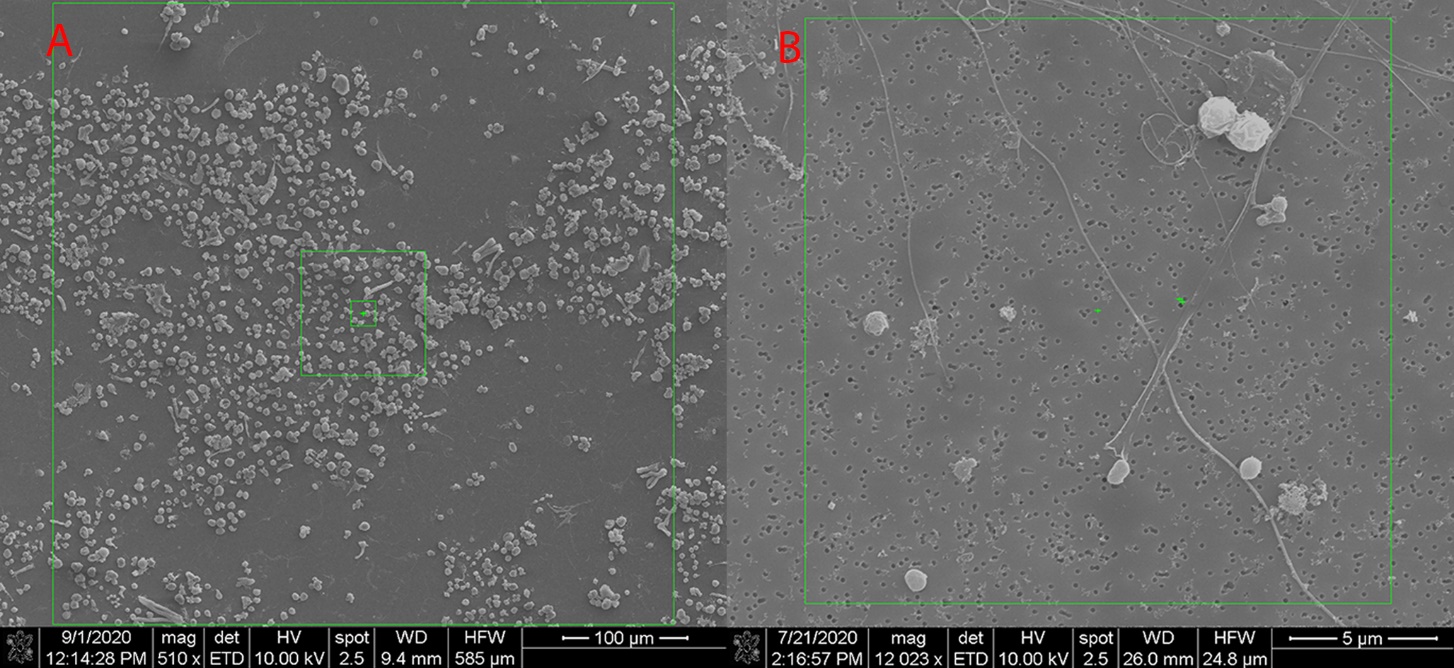

Supplementary Figure 3. Representative SEM images of A) unidentified white spherical particles and B) bacteria-like particles. Note the difference in scale as represented by the scale bar in the bottom right of each image.

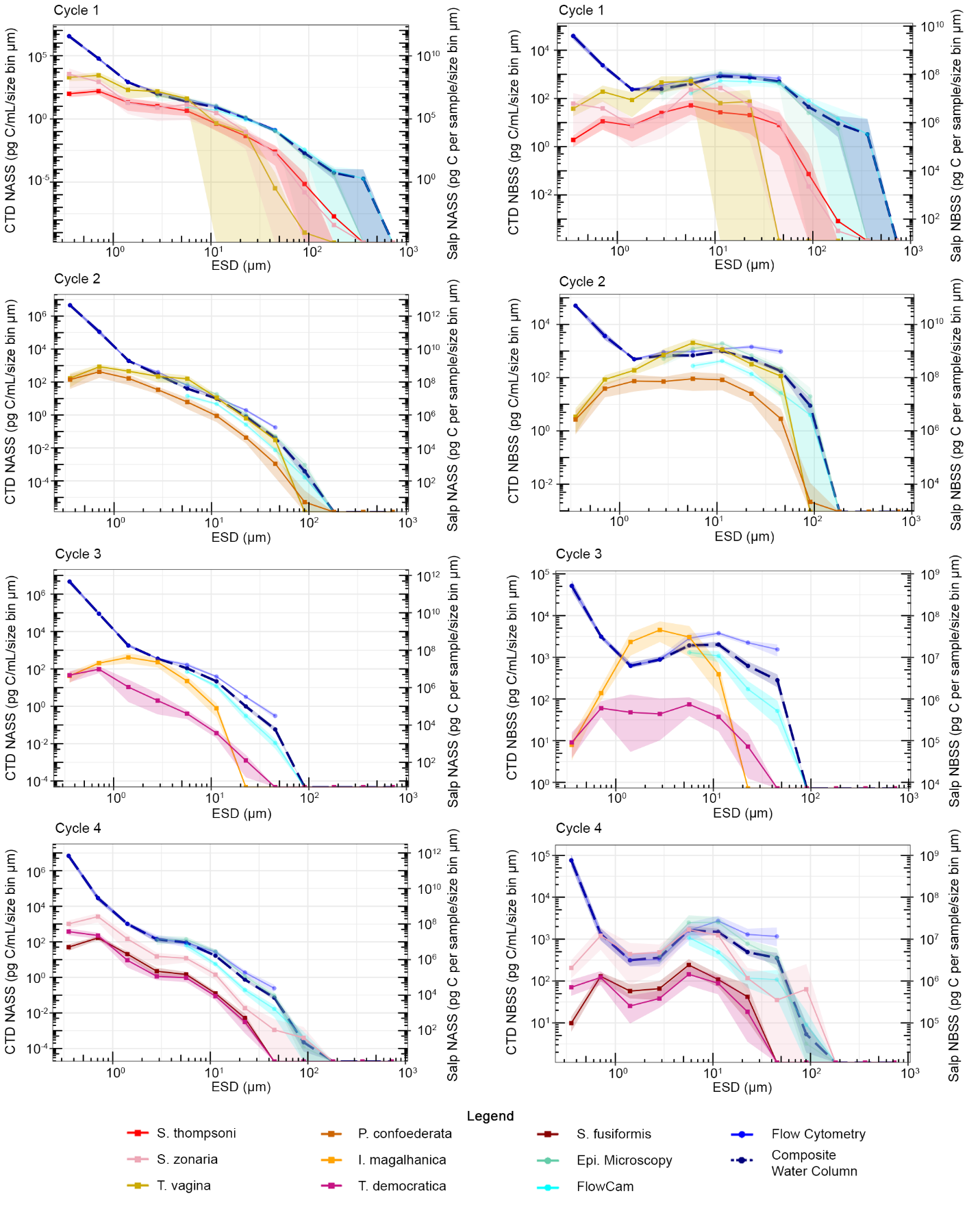

Supplementary Figure 4. Average normalized abundance (left) and normalized biomass (right) size spectra of salp prey (NASS and NBSS, respectively) as a function of equivalent spherical diameter (ESD) for Cycles 1-4. Lines and colors denote the same as Figure 3 in the main text, however all “broken” particles (less than ¾ assumed true ESD) are excluded.

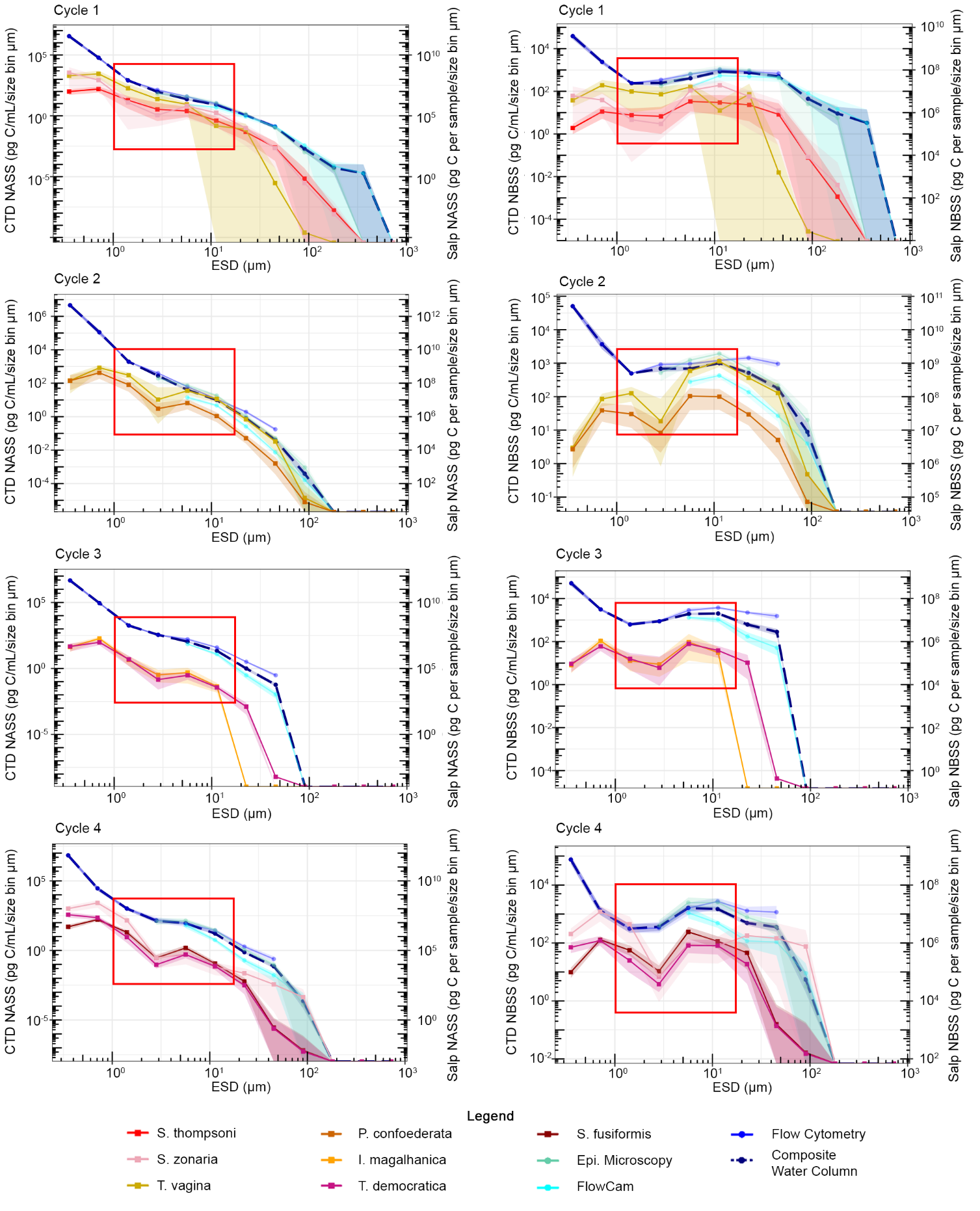

Supplementary Figure 5. Average normalized abundance (left) and normalized biomass (right) size spectra of salp prey (NASS and NBSS, respectively) as a function of equivalent spherical diameter (ESD) for Cycles 1-4. Lines and colors denote the same as Figure 3 in the main text, however “white sphere” particles are now removed and red boxes denote the size ranges over which they occurred.

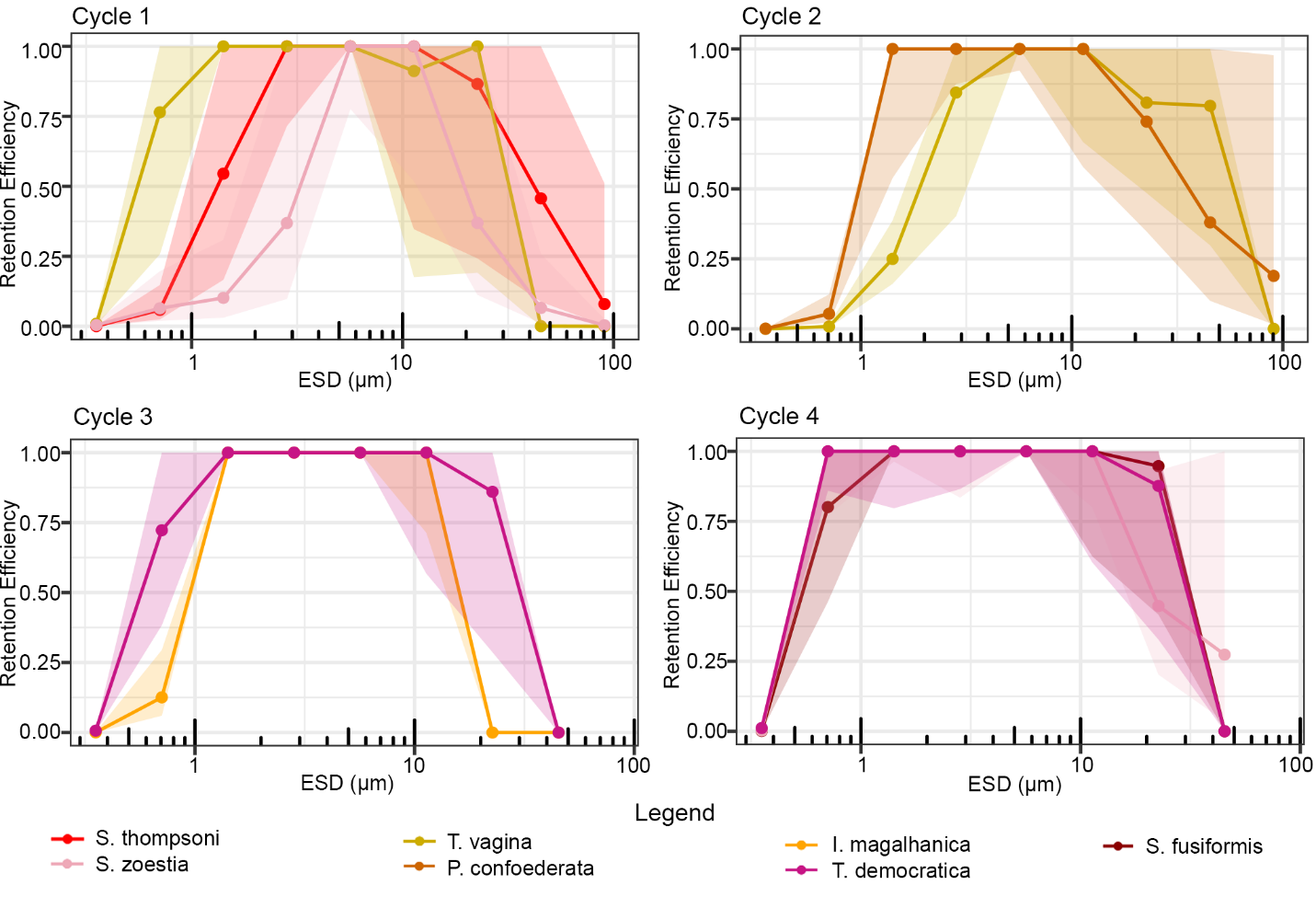

Supplementary Figure 6. Average retention efficiency as a function of prey equivalent spherical diameter (ESD) excluding broken particles for each salp species, organized by cycle, assuming filtration rate is equivalent to the clearance rate on 8-32 µm cells. Shaded areas represent 95% confidence intervals.

Supplementary Table 1. Salp community abundance (in individuals per 1000 m^-3^) determined via depth integrated MOCNESS catch for each salp species per cycle. Note that these do not correspond to the nets from which salps prepared for SEM were taken.

| **Species** | **Cycle** | **Abundance (ind 1000 m^-3^)** |
| --- | --- | --- |
| Salpa thompsoni | 1 | 1271.018126 |
| Thetys vagina | 1 | 10.29751569 |
| Soestia zonaria | 1 | 9.894001611 |
| Salpa thompsoni | 2 | 161.4036556 |
| Thetys vagina | 2 | 80.35189366 |
| Soestia zonaria | 2 | 3.966812567 |
| Pegea confederata | 2 | 15.2086029 |
| Ihlea magalhanica | 2 | 20.79745941 |
| Soestia zonaria | 3 | 4.710315591 |
| Ihlea magalhanica | 3 | 4.194630872 |
| Salpa fusiformis | 3 | 41.18965924 |
| Salpa thompsoni | 4 | 290.2357882 |
| Soestia zonaria | 4 | 18.92409938 |
| Pegea confederata | 4 | 35.8850674 |
| Salpa fusiformis | 4 | 9.957818891 |
| Thalia democratica | 4 | 136.5958765 |

Supplementary Table 2. Individual salp samples, the cycle from which they were collected, their species (IM = *Ihlea magalhanica*, SF = *Salpa fusiformis*, ST = *Salpa thompsoni*, TV = *Thetys vagina*, SZ = *Soestia zonaria*, PC = *Pegea confoederata*, TD = *Thalia democratica*), their life stage (A = aggregate, S = solitary), their total length after preservation, biomass-weighted mean prey size, minimum prey size, maximum prey size, log predator:prey size ratio (PPSR), and PPSR standard deviation (SD_PPSR_).

| **Sample** | **Cycle** | **Species** | **Life Stage** | **Length (mm)** | **BM Mean Prey Size (µm)** | **Min. Prey size (µm)** | **Max. Prey Size (µm)** | **log PPSR** | **SD_PPSR_** |
| --- | --- | --- | --- | --- | --- | --- | --- | --- | --- |
| 1 | 1 | ST | A | 10 | 2.88 | 0.45 | 8.16 | 3.54 | 0.25 |
| 2 | 1 | ST | A | 11 | 10.74 | 0.41 | 16.39 | 3.01 | 0.38 |
| 3 | 1 | ST | A | 11 | 3.04 | 0.41 | 10.60 | 3.56 | 0.35 |
| 4 | 1 | ST | A | 16 | 11.15 | 0.41 | 90.51 | 3.16 | 0.30 |
| 5 | 1 | ST | A | 18 | 33.57 | 0.41 | 181.02 | 2.73 | 0.15 |
| 6 | 1 | ST | A | 20 | 24.06 | 0.41 | 181.02 | 2.92 | 0.20 |
| 7 | 1 | ST | A | 22 | 19.95 | 0.40 | 181.02 | 3.04 | 0.36 |
| 8 | 1 | ST | A | 25 | 21.44 | 0.41 | 181.02 | 3.07 | 0.35 |
| 9 | 1 | ST | A | 66 | 9.93 | 0.41 | 181.02 | 3.82 | 0.25 |
| 10 | 1 | ST | A | 66 | 11.26 | 0.41 | 181.02 | 3.77 | 0.39 |
| 11 | 1 | ST | A | 68 | 8.46 | 0.41 | 181.02 | 3.91 | 0.41 |
| 12 | 1 | ST | S | 87 | 24.20 | 0.44 | 181.02 | 3.56 | 0.28 |
| 13 | 1 | ST | S | 104 | 5.43 | 0.42 | 13.20 | 4.28 | 0.15 |
| 14 | 1 | ST | S | 111 | 7.08 | 0.42 | 13.04 | 4.20 | 0.16 |
| 15 | 1 | SZ | S | 35 | 10.84 | 0.40 | 181.02 | 3.51 | 0.24 |
| 16 | 1 | SZ | S | 53 | 12.80 | 0.40 | 181.02 | 3.62 | 0.31 |
| 17 | 1 | SZ | S | 64 | 10.29 | 0.40 | 181.02 | 3.79 | 0.26 |
| 18 | 1 | TV | S | 143 | 7.42 | 0.41 | 90.51 | 4.29 | 0.35 |
| 19 | 2 | PC | A | 20 | 5.96 | 0.40 | 45.25 | 3.53 | 0.38 |
| 20 | 2 | PC | A | 20 | 10.72 | 0.40 | 45.25 | 3.27 | 0.30 |
| 21 | 2 | PC | A | 21 | 10.73 | 0.40 | 45.25 | 3.29 | 0.35 |
| 22 | 2 | PC | A | 22 | 10.54 | 0.41 | 45.25 | 3.32 | 0.26 |
| 23 | 2 | PC | A | 22 | 10.86 | 0.40 | 90.51 | 3.31 | 0.29 |
| 24 | 2 | PC | A | 38 | 11.45 | 0.42 | 45.25 | 3.52 | 0.33 |
| 25 | 2 | PC | A | 38 | 12.78 | 0.43 | 90.51 | 3.47 | 0.35 |
| 26 | 2 | PC | A | 45 | 11.37 | 0.40 | 93.17 | 3.60 | 0.45 |
| 27 | 2 | PC | A | 50 | 9.99 | 0.43 | 45.25 | 3.70 | 0.40 |
| 28 | 2 | PC | A | 72 | 12.25 | 0.41 | 90.51 | 3.77 | 0.37 |
| 29 | 2 | PC | A | 92 | 11.07 | 0.42 | 90.51 | 3.92 | 0.28 |
| 30 | 2 | PC | A | 95 | 10.21 | 0.41 | 90.51 | 3.97 | 0.28 |
| 31 | 2 | PC | A | 98 | 11.84 | 0.40 | 90.51 | 3.92 | 0.34 |
| 32 | 2 | PC | S | 30 | 3.56 | 0.41 | 45.25 | 3.93 | 0.23 |
| 33 | 2 | PC | S | 46 | 11.77 | 0.41 | 90.51 | 3.59 | 0.46 |
| 34 | 2 | PC | S | 62 | 10.95 | 0.40 | 90.51 | 3.75 | 0.46 |
| 35 | 2 | PC | S | 87 | 12.98 | 0.41 | 90.51 | 3.83 | 0.42 |
| 36 | 2 | PC | S | 126 | 11.57 | 0.41 | 90.51 | 4.04 | 0.34 |
| 37 | 2 | TV | S | 163 | 11.74 | 0.44 | 90.51 | 4.14 | 0.36 |
| 38 | 3 | IM | A | 16 | 4.53 | 0.41 | 11.26 | 3.55 | 0.24 |
| 39 | 3 | IM | A | 17 | 4.66 | 0.41 | 13.46 | 3.56 | 0.23 |
| 40 | 3 | IM | A | 17 | 3.81 | 0.42 | 9.18 | 3.65 | 0.28 |
| 41 | 3 | TD | A | 8 | 6.75 | 0.40 | 19.08 | 3.07 | 0.21 |
| 42 | 3 | TD | A | 9 | 6.20 | 0.41 | 11.51 | 3.16 | 0.43 |
| 43 | 3 | TD | A | 10 | 10.10 | 0.44 | 45.25 | 3.00 | 0.23 |
| 44 | 3 | TD | A | 12 | 7.10 | 0.41 | 12.78 | 3.23 | 0.28 |
| 45 | 3 | TD | A | 13 | 9.59 | 0.40 | 22.63 | 3.13 | 0.24 |
| 46 | 3 | TD | A | 14 | 7.33 | 0.41 | 45.25 | 3.28 | 0.45 |
| 47 | 4 | SF | A | 31 | 8.61 | 0.41 | 30.24 | 3.56 | 0.25 |
| 48 | 4 | SF | A | 33 | 7.86 | 0.42 | 22.63 | 3.62 | 0.24 |
| 49 | 4 | SF | A | 33 | 7.99 | 0.40 | 29.65 | 3.62 | 0.29 |
| 50 | 4 | SF | A | 35 | 6.81 | 0.40 | 90.51 | 3.71 | 0.21 |
| 51 | 4 | SF | A | 39 | 10.89 | 0.40 | 31.91 | 3.55 | 0.34 |
| 52 | 4 | SZ | A | 37 | 11.14 | 0.40 | 90.51 | 3.52 | 0.18 |
| 53 | 4 | SZ | A | 37 | 19.66 | 0.41 | 90.51 | 3.27 | 0.50 |
| 54 | 4 | SZ | A | 38 | 23.90 | 0.40 | 90.51 | 3.20 | 0.48 |
| 55 | 4 | TD | S | 14 | 7.18 | 0.40 | 22.63 | 3.29 | 0.17 |
| 56 | 4 | TD | S | 15 | 9.44 | 0.40 | 22.63 | 3.20 | 0.31 |
| 57 | 4 | TD | S | 15 | 12.01 | 0.40 | 30.57 | 3.10 | 0.31 |
| 58 | 4 | TD | S | 16 | 8.15 | 0.40 | 90.51 | 3.29 | 0.25 |
